# The Role of Genetic Variation in Shaping Phenotypic Responses to Diet in Aging *Drosophila melanogaster*

**DOI:** 10.1101/2025.01.09.632132

**Authors:** Nikolaj Klausholt Bak, Trudy F. C. Mackay, Fabio Morgante, Kåre Lehmann Nielsen, Jeppe Lund Nielsen, Torsten Nygaard Kristensen, Palle Duun Rohde

**Author notes:** These authors contributed equally (Torsten Nygaard Kristensen, Palle Duun Rohde).

## Abstract

Nutrition plays a central role in healthy living, however, extensive variability in individual responses to dietary interventions complicates our understanding of its effects. Here we present a comprehensive study utilizing the *Drosophila* Genetic Reference Panel (DGRP), investigating how genetic variation influences responses to diet and aging. Quantitative genetic analyses of the impact of dietary restriction on lifespan, locomotor activity, dry weight, and heat knockdown time were performed. Locomotor activity, dry weight and heat knockdown time were measured on the same individual flies. We found significant genotype-by-diet interaction (GDI) and genotype-by-age interaction (GAI) for all traits. Therefore, environmental factors play a crucial role in shaping trait variation at different ages and diets, and/or distinct genetic variation influences these traits at different ages and diets. Our genome wide association study also identified a quantitative trait locus for age-dependent dietary response. The observed GDI and GAI indicates that susceptibility to environmental influences changes as organisms age, which could have significant implications for dietary recommendations and interventions aimed at promoting healthy aging in humans. The identification of associations between DNA sequence variation and age-dependent dietary responses opens new avenues for research into the genetic mechanisms underlying these interactions.

## Introduction

All organisms age and eventually die – a topic that has interested researchers for decades (1–3). Lifespan and healthspan are defined as the total time an individual lives and the period of life during which individuals are healthy and free from age-related diseases, respectively (4–6). Extending lifespan without improving healthspan can lead to longer periods of poor health and diminished quality of life (5,7). In humans and model species such as mice (*Mus musculus*), roundworms (*Caenorhabditis elegans*), and vinegar flies (*Drosophila melanogaster*), it is well established that lifespan and healthspan can be uncoupled (6–10). This means that the duration of healthy life can be altered without necessarily changing the overall lifespan, and *vice versa*. Therefore, living longer does not always mean better health late in life, highlighting the importance of studying both lifespan and healthspan to uncover mechanisms affecting each (6,11). Obviously, this topic is of general importance and interest to humans, as the average global lifespan has increased markedly by ca. 21 years from 1960 to 2020 (12). However, healthspan has not increased at the same pace (13–15). This discrepancy poses significant societal challenges because it reduces the quality of life for the elderly and imposes economic burdens on healthcare systems. However, the interest in aging extends beyond humans. For livestock, improving healthspan can enhance animal welfare. Livestock are raised for economic purposes and are often culled when their production declines. By improving the overall quality of life and health of these production animals, it may be possible to extend their productive years. This approach could reduce costs for farmers and enhance the welfare of the animals (16). In wild populations, understanding aging can help in conservation efforts (17,18), and in evolutionary biology studying aging can provide insights into the natural selection processes that shape lifespan and healthspan within and across different species (19–21).

The considerable variation in age-related traits raises fundamental questions: Why does the average lifespan differ so much between even closely related species? How do we explain why Greenlandic sharks (*Somniosus microcephalus*) can live 500 years and that closely related species such as the Pacific sleeper (*S. pacificus*) ‘only’ live up to 250 years (22)? Why do some genotypes of *D. melanogaster* have an average lifespan of 22 days, while others survive up to 80 days (23). Additionally, *D. melanogaster* and *C. elegans* individuals of the same genotype reared under different environmental conditions can vary in lifespan by more than a factor of 5 (24,25). These variations arise due to genetic and environmental factors, as well as interactions between genotype and environment.

To address these questions on the genetic basis of variation in lifespan and healthspan across age classes and nutritional conditions, we utilized a subset of the *D. melanogaster* Genetic Reference Panel (DGRP), which includes more than 200 inbred lines with full genome sequences (26). Using the DGRP in genetic studies provides many advantages, including the rapid decay of linkage disequilibrium in *D. melanogaster* which allows for precise mapping of genetic variants (26). The DGRP system allows testing many individuals with the same genotype in different environments, enabling an understanding of the interaction between genetic variation and environmental conditions (26–31).

The DGRP has been instrumental in identifying genes that affect various traits, including aging and response to diets, providing potential targets for anti-aging interventions (11,25,32). Healthspan studies in *D. melanogaster* typically use measures such as locomotor activity, stress responses, and body mass as health indicators. Locomotor activity in flies often declines with increased age or disease, which makes it a valuable healthspan metric (11,33). Rapid iterative negative geotaxis and *Drosophila* activity monitor (DAM) assays measure locomotor ability and overall activity, respectively, while body size indicates fitness influenced by environmental factors (34–36). Heat stress resistance is another important metric, as higher resistance often correlates with longer lifespan, better overall health and resilience (37–40).

Here, we investigated the effects of a standard nutritionally rich diet and a restricted diet on lifespan and healthspan metrics in *D. melanogaster*; specifically, locomotor activity, heat knockdown time (HKDT), and dry weight. Our study aimed to investigate whether (i) there is genetic variation for all traits across diet environments; (ii) assess genotype-by-diet and genotype-by-age interactions for lifespan, dry weight, locomotor activity and HKDT; (iii) identify candidate genes associated with environment-specific or pleiotropic allelic effects within the same trait using genome-wide studies (GWAS).

## Materials and Methods

### Fly Stocks and Nutritional Environments

We used 98 DGRP lines, obtained from the Bloomington *Drosophila* Stock Center (NIH P40OD018537). The DGRP is a collection of inbred lines derived from a natural population of *D. melanogaster* from Raleigh, North Carolina, USA. Each line has undergone 20 generations of full-sib inbreeding, resulting in extremely low genetic variation within lines and genetic variation reflecting the original population distributed between lines (41,42). The complete genomes of all DGRP lines have been sequenced to high coverage (average ∼27 fold), identifying a total of 4,565,215 molecular variants, including single or multiple nucleotide polymorphisms, insertions, deletions, and microsatellites.

Prior to the experiments, flies were maintained in a climate-controlled room at 23°C and 50% relative humidity with a 12:12 h light/dark cycle. Flies were reared on either a standard or restricted diet. The standard diet was Leeds medium composed of dry yeast (60 g·L^-1^), sucrose (40 g·L^-1^), oatmeal (30 g·L^-1^), agar (18 g·L^-1^), Nipagen (12 mL·L^-1^) and acetic acid (1.2 mL·L^-1^). The restricted diet was a 25% dilution of the nutritional content of the standard Leeds medium using the indigestible compound α-cellulose (Product no. 102550125, Sigma-Aldrich, Buchs SG, Switzerland) (Table 1). The concentrations of agar, nipagin and acetic acid were the same in the two diet types. This restricted diet has previously been shown to cause marked decreases in traits related to lifespan and healthspan (8).

**Table 1.**
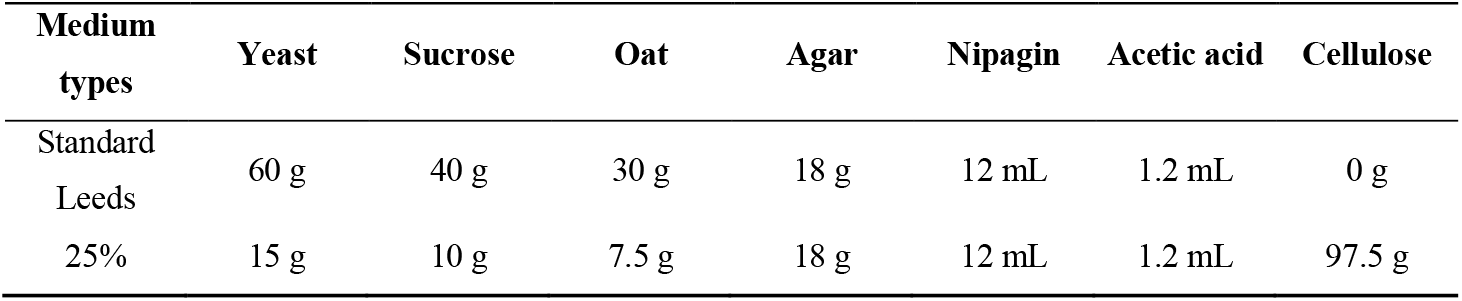
Nutritional components of the two different medium types. The control diet is set to have 100% nutritional value and the nutritional value of the restrictive diet is set relative to that.

### Experimental Design

For each DGRP line, ca. 10 flies were transferred to new vials every third day 5-7 times, resulting in 15-20 vials containing the F1 generation. The F1 generation flies were pooled into five bottles per line, each containing 100 flies, to ensure consistent density, provided there were enough flies. The flies were transferred to new bottles daily for a total of five days, resulting in up to 25 bottles per line. Upon emergence of the F2 generation, males were retained while females were discarded. Males were used in the experiments because it was not feasible to ensure that the females were virgins, and females have a higher reproduction and age trade-off than males (43–45). This sorting procedure was continued for four days, and on the fifth day, the flies were 2 days ± 36 hours old when placed into three bottles (whenever possible) for each diet type, with each bottle containing exactly 110 flies (Figure 1). This resulted in a total of 6,338 flies, with 4,015 flies reared on the control diet, and 2,323 flies reared on the restricted diet for the measurement of the healthspan metrics; dry weight, locomotor activity and heat knockdown time (HKDT). These health metrics were also measured on 7 day-old flies and at 9 day intervals until flies were 61 day-old (N = 1,561 at test day 7; N = 1,504 at test day 16; N = 1,440 at test day 25; N = 1,126 at test day 34; N = 530 at test day 43; N = 170 at test day 52 and N = 7 at test day 61). Generally, flies had a shorter lifespan when fed on the restricted diet compared to those fed on the control diet, which resulted in an unbalanced data set. The healthspan dataset was balanced by removing lines where all the flies had died on one diet (for the number of flies on each diet and age see Tables S1-S3). Analyses run with all data included gave similar results, but we had greater power for all health metrics when working with the balanced dataset (Figure S1 and Tables S4-S9). With increasing age, fewer individuals were alive for phenotyping, leading to a significant reduction in statistical power. Consequently, data from day 25 and beyond were removed, and we present here the results from analysis of data from days 7 and 16.

**Figure 1.**
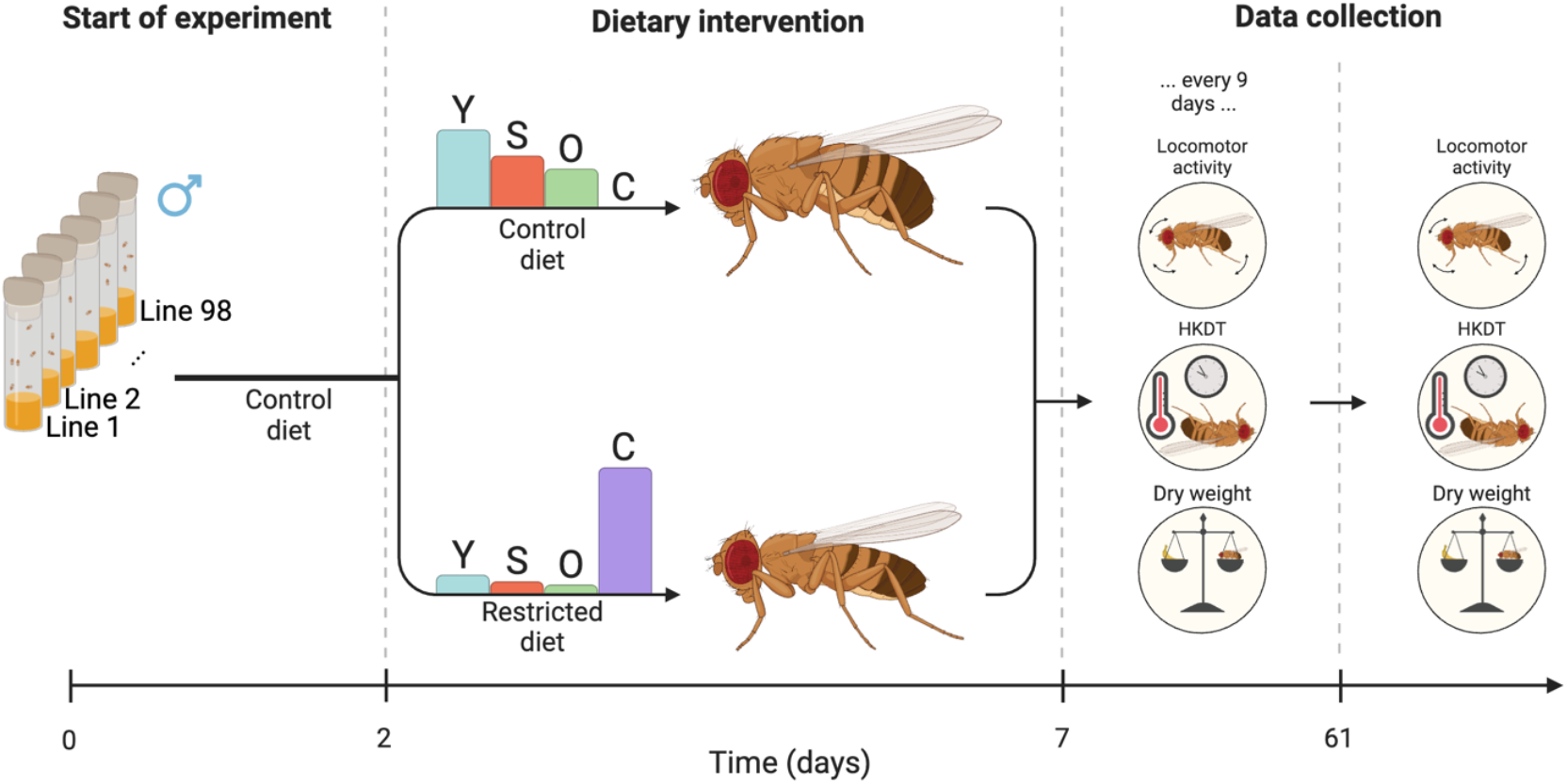
Flowchart of experiment. 98 DGRP lines were raised at 23°C on a control diet. When the adult flies were two days ± 36 hours old, they were placed on either a control diet (Leeds medium) or a restricted diet. Health metrics, including locomotor activity, HKDT, and dry weight, were first assessed when the flies were seven day-old and subsequently measured every nine days. From the same pool of flies, mortality was recorded every three days, starting from five day-old flies. Barplots indicate the relative content of the dietary components yeast (Y), sugar (S), oat (O) and cellulose (C).

### Lifespan Assay

For each DGRP line and dietary condition (Table 1), we attempted to rear three biological replicates with 110 flies each. This was not possible for all lines. The cohorts were transferred to fresh vials containing 60 mL of medium every three days. During these transfers, mortality was recorded, and the deceased individuals were removed from the study. This protocol was followed until all flies had died in all replicates. Lifespan was measured on a total of 35,067 flies (15,904 on the control diet (on average 166 flies/line) and 19,163 on the restricted diet (on average 196 flies/line).

### Locomotor Activity Assay

For the first locomotor activity assay, eight seven day-old flies from all three replicate bottles per line and nutritional condition were transferred to 5 mm polycarbon tubes (TriKinetics Inc, Waltham, MA USA) containing a pipe cleaner moistened in water in both ends. We aimed to test 16 flies per line and nutritional condition for all other locomotor activity assays. However, flies from some lines survived longer than others and therefore fewer individuals were used at the later activity measurements for short-lived lines (Tables S1-S3). The polycarbon tubes containing flies were added to *Drosophila* Activity Monitors (DAM) (DAM2 Activity Monitor, Trikinetics Inc, Waltham, MA US). The monitors were placed in a climate-controlled chamber (Binder KB 400, Binder, Tuttlingen, Germany) at 23°C. Locomotor activity was quantified as the number of times individual flies generate an activity count every 10 seconds for 6 hours (9:40 PM to 3:40 AM) (Figure 2). Temperature and humidity conditions were recorded every 5 minutes.

**Figure 2.**
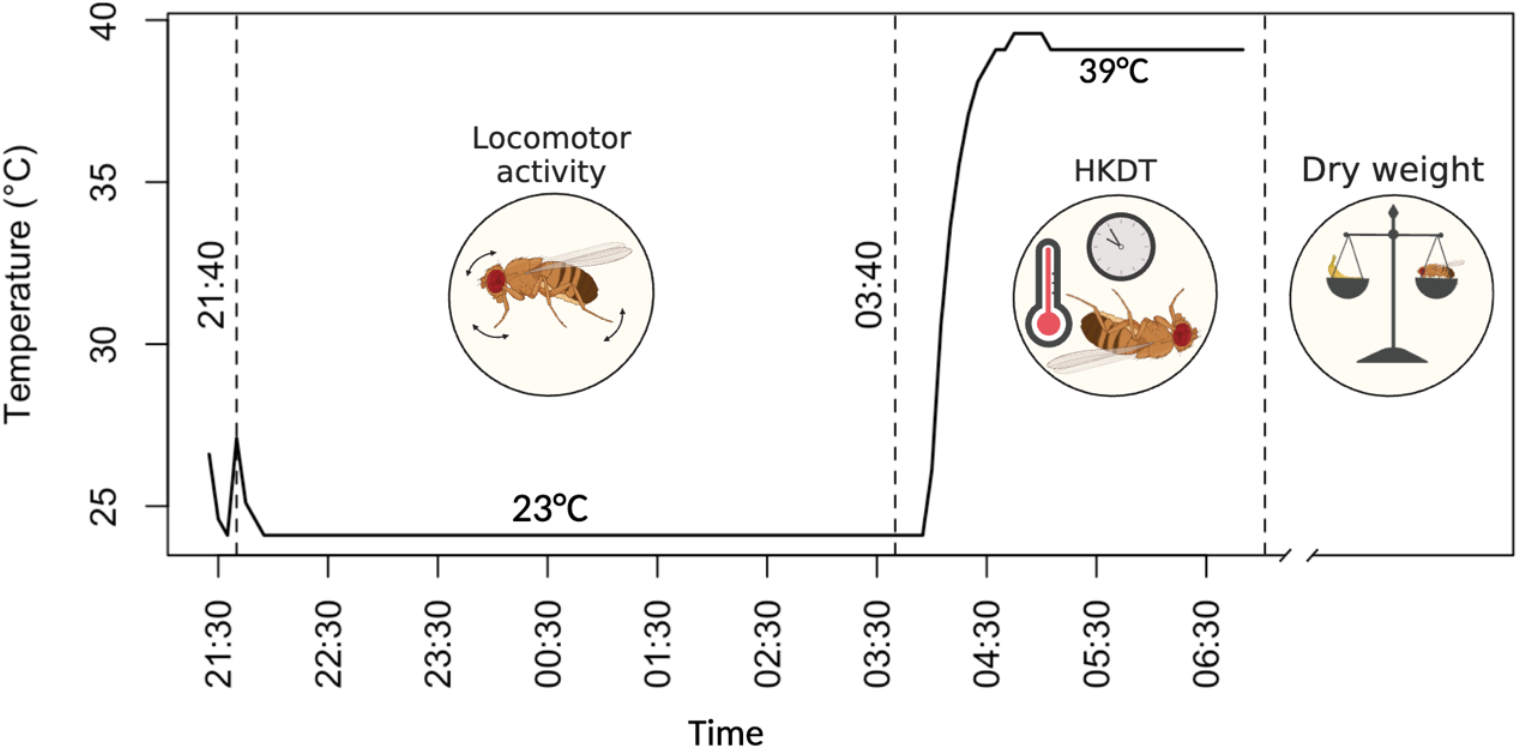
Flowchart of healthspan metrics (locomotor activity, heat stress tolerance and dry weight). Healthspan metrics were all obtained on the same individual flies. First, locomotor activity was monitored for six hours at 23°C. Subsequently, the HKDT assay started with a temperature increase to 39°C and HKDT was determined as the last recorded activity count using DAM. Finally, the flies were stored at -80°C for subsequent measurement of dry weight. The x-axis indicates the time of day.

### Heat Stress Tolerance

Following the assessment of locomotor activity at 23°C, the flies were kept in the activity monitors and exposed to 39°C within a climate-controlled chamber (Binder KB 400, Binder, Tuttlingen, Germany). Locomotor activity was recorded every 10 seconds for 2 hours (3:40 AM to 5:40 AM), after which all flies were dead (Figure 2). The time to death due to heat stress was determined as the last recorded activity count. Temperature and humidity conditions were monitored every 5 minutes. This assay provided data on HKDT. Individual flies used for locomotor activity and heat tolerance assessments were stored in Eppendorf tubes at -80°C for subsequent dry weight measurement.

### Dry Weight

After assessing heat stress tolerance, the flies were desiccated for 48 hours at 60°C and subsequently weighed to the nearest 10 µg (Sartorius Quintix35-1S, Satorius, Göttingen, Germany).

### Quantitative Genetic Analyses

The contributions of age, diet and DGRP line to variation in lifespan, locomotor activity, HKDT and dry weight in the DGRP were assessed separately using mixed model analyses of variance (ANOVA) with a type 3 estimation method utilizing PROC MIXED in SAS (Version 5.2, SAS Institute, Cary, NC). The full mixed model is shown in Eq. 1:@

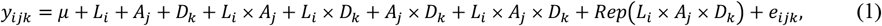

where *y*_*ijk*_ is the phenotype; *µ* is the fixed effect of the overall mean; *L*_*i*_ (*i* = 1, 2, …, 98) is the random effect of DGRP line; *A*_*j*_ (*j* = 1, 2, 3) and *D*_*K*_ (*k* = 1, 2) are the fixed effects of age and diet respectively; *L*_*i*_×*A*_*j*_, *L*_*i*_ × *D*_*k*_, *A*_*j*_×*D*_*k*_ and *L*_*i*_×*A*_*j*_ × *D*_*k*_ are the interaction effects; *Rep* is the random effect of replicate; and *ε*_*ijk*_ is the random residual effect. Broad-sense heritability, *H*^2^, for the full model was estimated as:

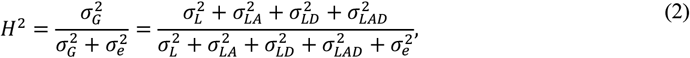

where 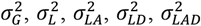 and 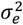 are the genetic, among-line, line × age, line × diet, line × age × diet and residual (environmental) variance components, respectively. All combinations of reduced ANOVA models were performed; all ages on separate diets, separate ages on all diets, and separate ages on all separate diets. Standard errors (SE) and confidence intervals (CI) of the *H*^2^ were calculated as in Rohde et al. (46):

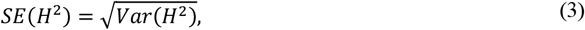

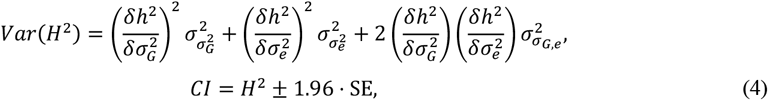

where 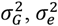, and 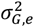 are the elements from the asymptotic covariance matrix. Moreover, the partial derivatives are 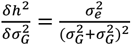, and 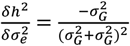. Genetic correlations, 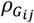, between ages and diets were estimated as Falconer and Mackay (47):

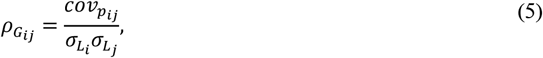

where 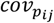 is the phenotypic covariance between a trait (*i*) at one diet and age (7 or 16 days) and trait (*j*) at the same or another diet or age. The percent contribution of changes in rank order (% rank), was calculated as in Cockerham (48):

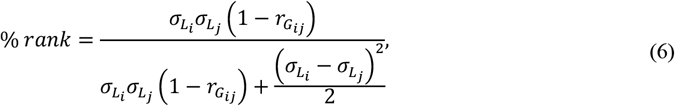

where 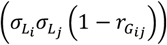 represents the component of the genotype-by-diet interaction (GDI) or genotype-by-age interaction (GAI) caused by changes in rank order of DGRP line means between the control and restricted diets or between ages 7 and 16 days. 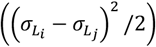 is the component of GDI or GAI attributed to differences in the magnitude of between-line genetic variance between the control and restricted diets or ages 7 and 16 days.

Phenotypic correlations were estimated using Pearson correlation coefficients (*ρ*) (equation 7):

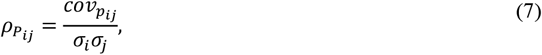

where *σ* is the trait value between a trait (*i*) at one diet and age (7 or 16 days) and trait (*j*) at the same or another diet or age. Standard errors and confidence intervals of the phenotypic and genotypic correlations were estimated as in Gnambs (49):

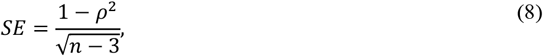

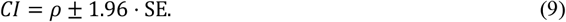

### Survival Analysis

The contribution of diet to the age of the DGRP lines was also assessed using a Cox Proportional Hazards model. The risk of events over time, *λ*(*t*), was estimated by:

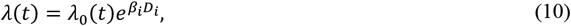

where *λ*_0_(*t*) is the baseline hazard, *D*_*i*_ is diet with *i* representing control or restricted diet, and *β*_*i*_ is the regression coefficient.

### Genome-wide Association Study

Single marker regression analyses were conducted on DGRP line means for lifespan, locomotor activity, HKDT and dry weight separately. We performed genome-wide association studies (GWAS) separately for diets and ages using a mixed model accounting for *Wolbachia* infection status, major chromosomal inversion status (*In(2L)t, In(2R)NS, In(3R)P, In(3R)K* and *In(3R)Mo*), and polygenic relatedness as described previously (25,41,42)). These analyses assessed the strength of association for additive effects of 1,928,067 polymorphic variants across 98 lines with a minor allele frequency (MAF) greater than 0.05. These analyses were performed for traits with significant genetic components (*p <* 0.05) in the ANOVA models.

We conducted GWAS for line means within each age group, considering two pairs of environments (control diet *vs*. restricted diet) separately for 7 and 16 day-old flies. Additionally, genotype-by-diet and genotype-by-age interaction effects were investigated using the difference in line means between the conditions as the analysis variable.

## Results

We assessed how genetic variation impacts responses to diet and aging using the *Drosophila* Genetic Reference Panel (DGRP). We performed quantitative genetic analyses to evaluate the effects of dietary restriction on lifespan, locomotor activity, dry weight, and heat knockdown time. These measurements were all taken from the same individual flies.

Dry weight ranged from 0.07 mg to 0.36 mg, with a mean of 0.18 mg (Figures 3A and S2A-B, Tables 2 and S10). A significant difference was observed between the control diet and restricted diet for 7 and 16 day-old flies (*p* < 0.0001; Table S7), with flies on the control diet having, on average, a 7% (0.01 mg) higher dry weight than flies on the restricted diet. Dry weight varied between lines for combinations of individual diets and ages (Table S7). There was no significant change in dry weight as the flies aged from 7 to 16 days on control diet (*p* = 0.4801, Table S7) or on restricted diet (*p* = 0.2610, Table S7), but there was an interaction effect between line and age on these diets (control diet; *p* < 0.0001, restricted diet; *p* < 0.0001, Table S7). Similar to dry weight, there was a significant decrease in lifespan when flies were exposed to a restricted diet compared to a control diet (Figures 3D, S2G-H and S3, and Tables 2, S11, S12 and S13). In contrast, locomotor activity was higher for flies exposed to the restricted diet (Figures 3B and S2C-D, Tables 2, S8 and S14), while HKDT is age-dependent (Figures 3C and S3E-F, Tables 2, S9 and S15). Locomotor activity plasticity with diet is age-dependent, where only 16 day-old flies showed a significant difference in locomotor activity between the two diets. Furthermore, locomotor activity differs across age in a diet-dependent manner, where only flies kept at the control diet showed plasticity with regards to age. HKDT showed plasticity for both diet and age. Similarly, lifespan showed diet-dependent plasticity.

**Figure 3.**
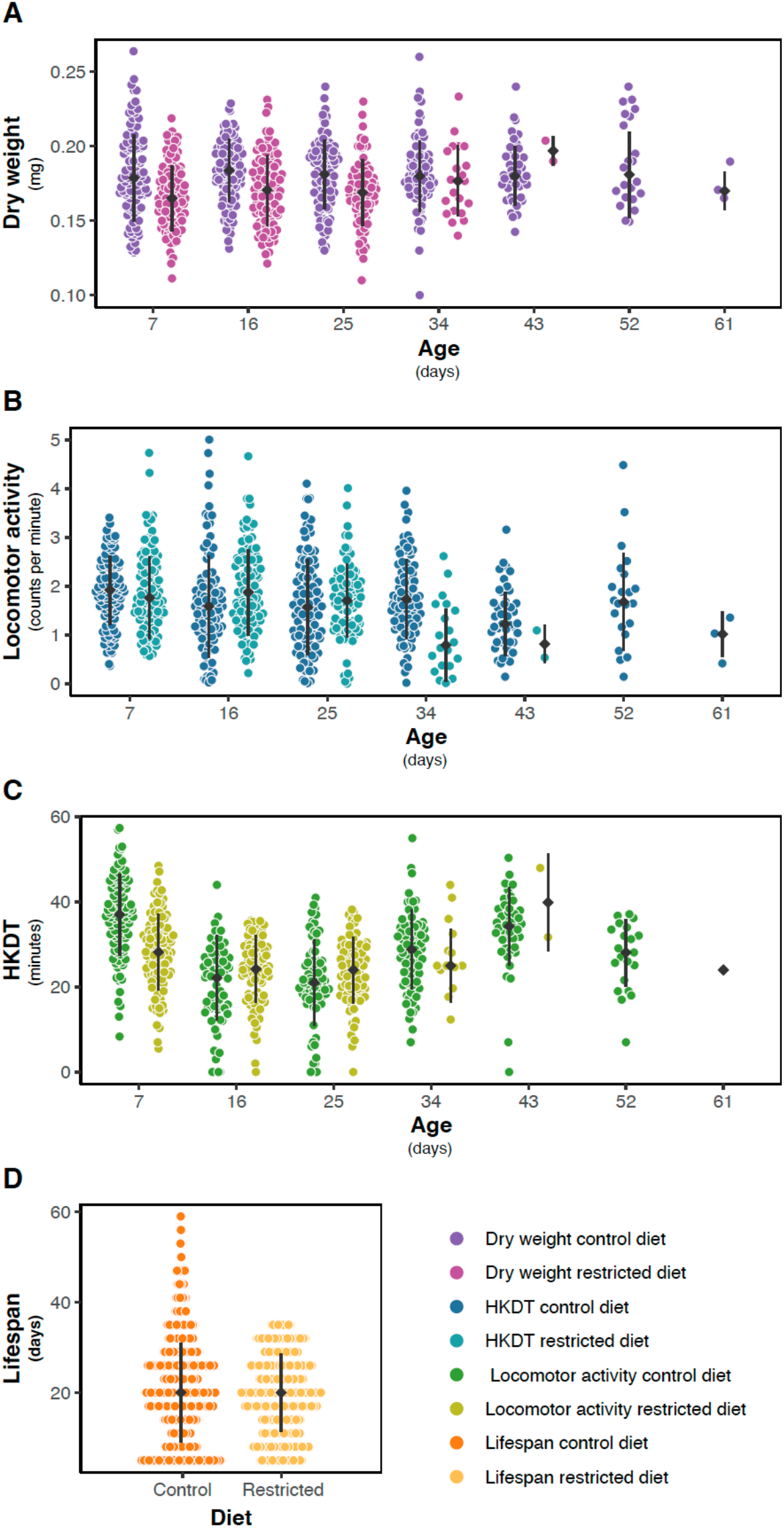
Distribution of line means. Mean (**A**) dry weight (purple), (**B**) locomotor activity (blue) and (**C**) HKDT (green) of DGRP lines across seven ages, fed on control diet (dark shades) and restricted diet (light shades), and (**D**) lifespan (orange) also fed on control diet (dark shade) and restricted diet (light shade). Points represent the mean phenotype of each DGRP line, with filled grey points indicating the mean population trati values and grey lines indicating population standard errors for each diet and age.

**Table 2.** *A*verage trait means within diet and age groups at the two different diets.

| Phenotype | Control diet |  | Restricted diet |  |
| --- | --- | --- | --- | --- |
|  | 7 day-old | 16 day-old | 7 day-old | 16 day-old |
| Dry weight | 0.182 | 0.183 | 0.168 | 0.171 |
| Locomotor activity | 1.890 | 1.869 | 1.918 | 2.022 |
| HKDT | 37.11 | 22.38 | 29.91 | 25.40 |
| Lifespan | 20.58 |  | 19.34 |  |

To understand the relationships between the various traits, age groups, and diets, we estimated the phenotypic correlations among them. Strong positive correlations between the same traits across different age groups and diet types were observed for all traits (Figure 4 and Tables S16-S17). Most of the phenotypic correlations between traits were positive, with the exception of locomotor activity in 7 day-old flies fed the control diet and HKDT in 16 day-old flies fed the control diet, which showed negative correlations with other traits (Figure 4 and Tables S16-S17).

**Figure 4.**
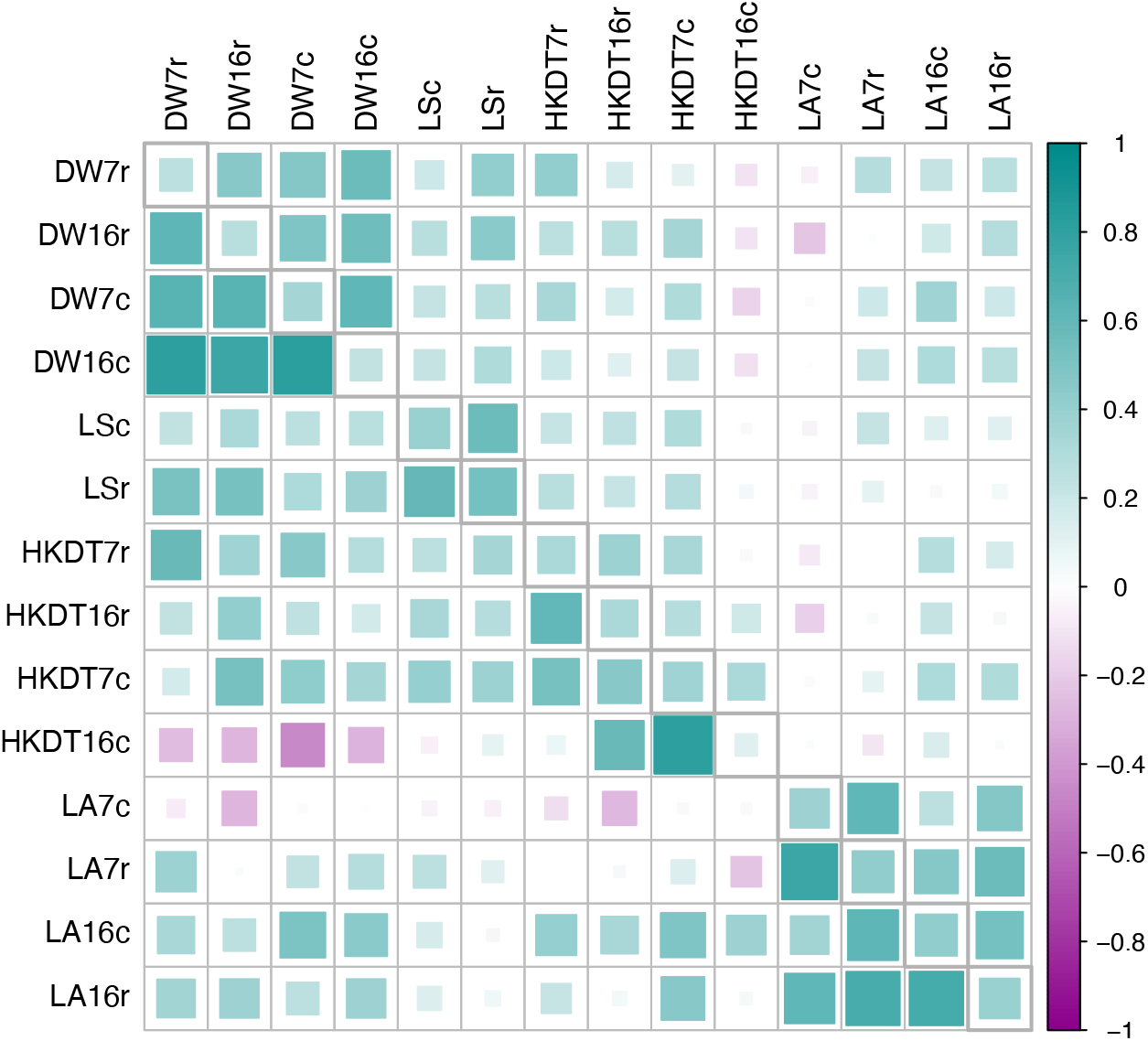
Phenotypic (above diagonal) and genetic (below diagonal) correlations with heritability estimates in the diagonal for all traits. This figure illustrates the linear relationships between traits, age groups, and diet types. The traits are dry weight (DW), lifespan (LS), heat knockdown time (HKDT), and locomotor activity (LA). Age groups are 7 day-old (7) and 16 day-old (16) flies, and diet types are control diet (c) and restricted diet (r). The upper diagonal matrix displays the phenotypic Pearson correlation coefficients between traits, the diagonal contains the estimated heritabilities and the lower diagonal matrix shows the genetic correlations between traits. Color and size indicate the strength of the correlations and the heritability (cyan for positive correlations and magenta for negative correlations, with a larger size indicating a stronger correlation).

The estimates of broad sense heritability 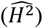 for dry weight ranged from 0.24 to 0.34, for locomotor activity from 0.37 to 0.43, for HKDT from 0.12 to 0.36 and for lifespan from 0.39 to 0.53 (Figure 4 and Tables S16-S17). Most of the estimated environmental variance components, genetic variances components and heritabilities were significantly different from zero, but not significantly different between diets and ages within traits (Figure 4 and Tables S16-S17).

The cross-diet genetic correlation for dry weight between control and restricted diets 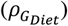 was 0.645 at 7 days and 0.755 at 16 days (Figure 4 and Tables S16-17). The interaction effect between diet and line was significant at both day 7 (*p* < 0.0001, Table S7) and day 16 (*p* = 0.0030, Table S7). Therefore, the genetic correlations at both ages were signficantly less than unity, indicating genotype-by-diet interaction (GDI) for this trait (Figure 5). The GDI variance was primarily due to changes in rank order of line means of dry weight rather than between-line variance for the two diets within each age group. The GDI variance for dry weight was increasingly attributed to the rank order, from 73.5% at day 7 to 96.8% at day 16 (Figure 6 and Table S18). The cross-age genetic correlation for dry weight of 7 and 16 day-old flies 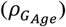 was higher on the control diet (0.823) than the restricted diet (0.623) (Figure 4 and Tables S16-17). Since the interaction effect between age and line was significant both on the control diet (*p <* 0.0001, Table S7) and on the restricted diet (*p =* 0.0003, Table S7), the correlations on both diets were significantly less than unity, indicating genotype by age interaction (GAI) for this trait (Figure S4). The GAI variance was mostly due to differences in the rank order of line means of dry weight between day 7 and day 16 on the control diet (93.6% rank, Figure 6 and Table S18) and on the restricted diet (99.4% rank, Figure 6 and Table S18). Locomotor activity, HKDT, and lifespan exhibited highly significant interaction effects between both diet and line and age and line, indicating GDI and GAI for these traits as well (Tables S8-S10). The GDI was primarily influenced by the rank order of line means for locomotor activity, HKDT and lifespan, which ranged from 88.8% to 100% (Figure 6 and Table S18). GAI was also largely due to the rank order of line means for locomotor activity and HKDT with values increasing from 84.3% to 100% (Figure 6 and Table S18).

**Figure 5.**
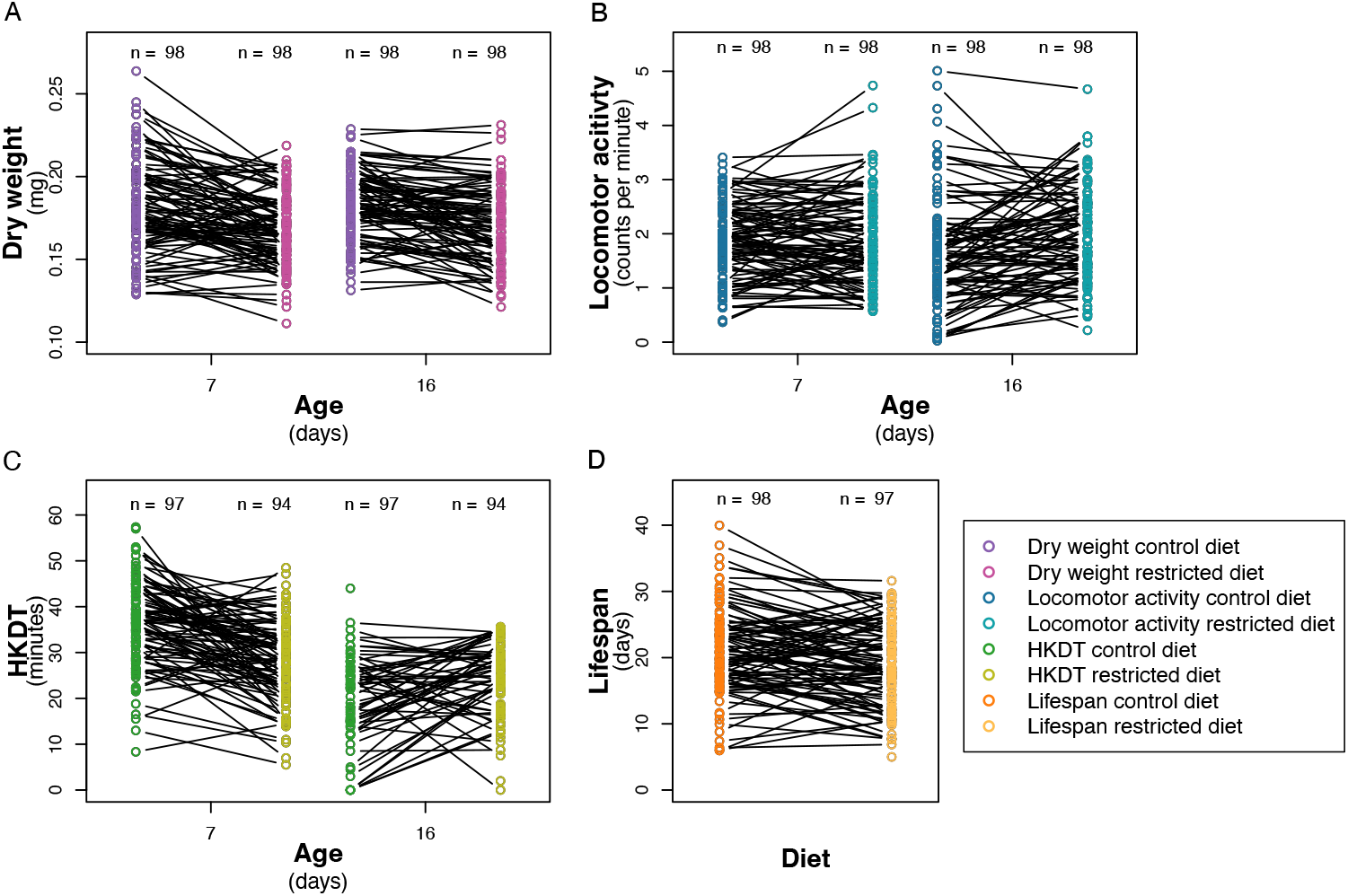
Reaction norm plot. Line means of (**A**) dry weight (purple), (**B**) locomotor activity (blue) and (**C**) HKDT (green) across seven ages, fed on control diet (dark shades) and restricted diet (light shades), and (**D**) lifespan (orange) also fed on control diet (dark shades) and restricted diet (light shades). Points represent the mean phenotype of each DGRP line, and lines connecting points across diets represent the same DGRP line. Numbers (n) of DGRP lines assessed at each age and diet is indicated above the points.

**Figure 6.**
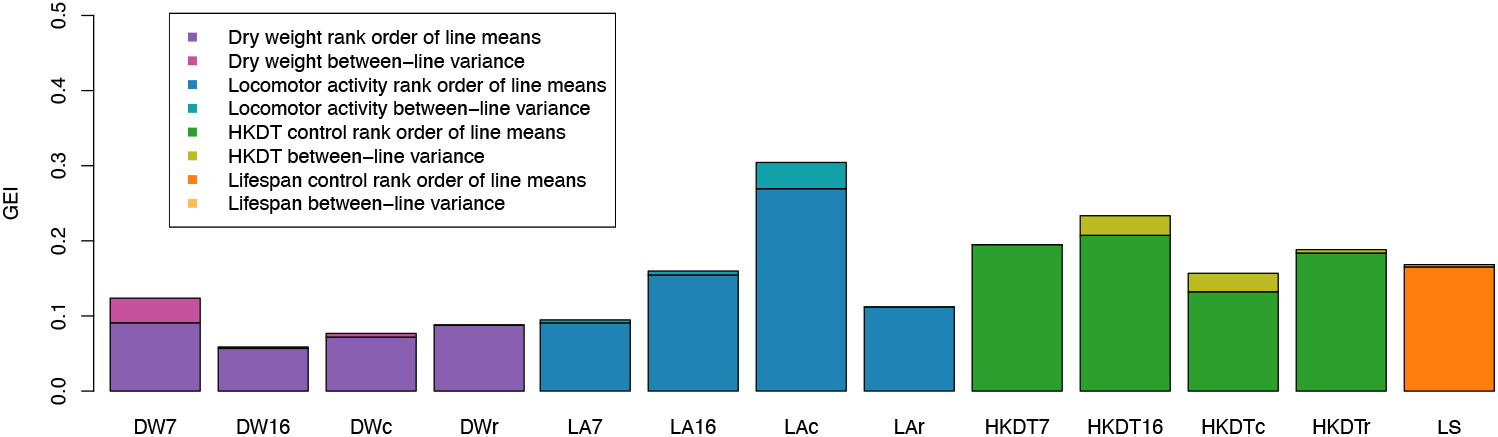
GAI and GDI due to differences in magnitude of between line variance and changes in rank order. The relative proportion of interaction variance due to these effects for genotype-by-diet or genotype-by-age interactions for dry weight (DW), lifespan (LS), heat knockdown time (HKDT), and locomotor activity (LA). The age groups are 7 day-old (7) and 16 day-old (16) flies, and the diet types are control (c) and restricted (r) diets. Colors indicate the proportion of GDI and GAI attributed by differences in magnitude of between-line variance (light shades) of genetic variance between the control and restricted diet or age 7 and 16 days and rank order of line means (dark shades).

We performed GWAS for dry weight, locomotor activity, HKDT, and lifespan using 1,928,067 polymorphic (minor allele frequency > 0.05) genetic variants in 98 DGRP lines. We conducted GWAS using line means within each age group, considering two pairs of environments (control diet *vs*. restricted diet) separately for 7- and 16 day-old flies. Additionally, diet-by-line and age-by-line interaction effects were investigated by performing the GWAS using the difference in line means between conditions.

Genotype-by-diet interaction for dry weight showed a strong signal for the variants in the gene encoding Peptidoglycan Recognition Protein LC (*PGRP-LC*), as indicated by the Quantile-Quantile (Q-Q) plot (Figure 7A) and -log_10_(*p*) values below the Bonferroni threshold in the Manhattan plot (Figure 7B and Table S19) *PGRP-LC* is essential for the innate immune response in *D. melanogaster* (50,51). No other genetic variants were associated with dry weight, locomotor activity and HKDT (Figures S5-S7) association tests at the genome-wide significance threshold. However, a single genetic variant within the gene muscleblind (*mbl*) was genome-wide significant for the lifespan genotype-by-diet interaction (Figures 7C-D and Table S20). *mbl* encodes an RNA-binding protein that plays a crucial role in RNA metabolism, including alternative splicing, transcript localization, and the biogenesis of miRNA and circRNA (52–54).

**Figure 7.**
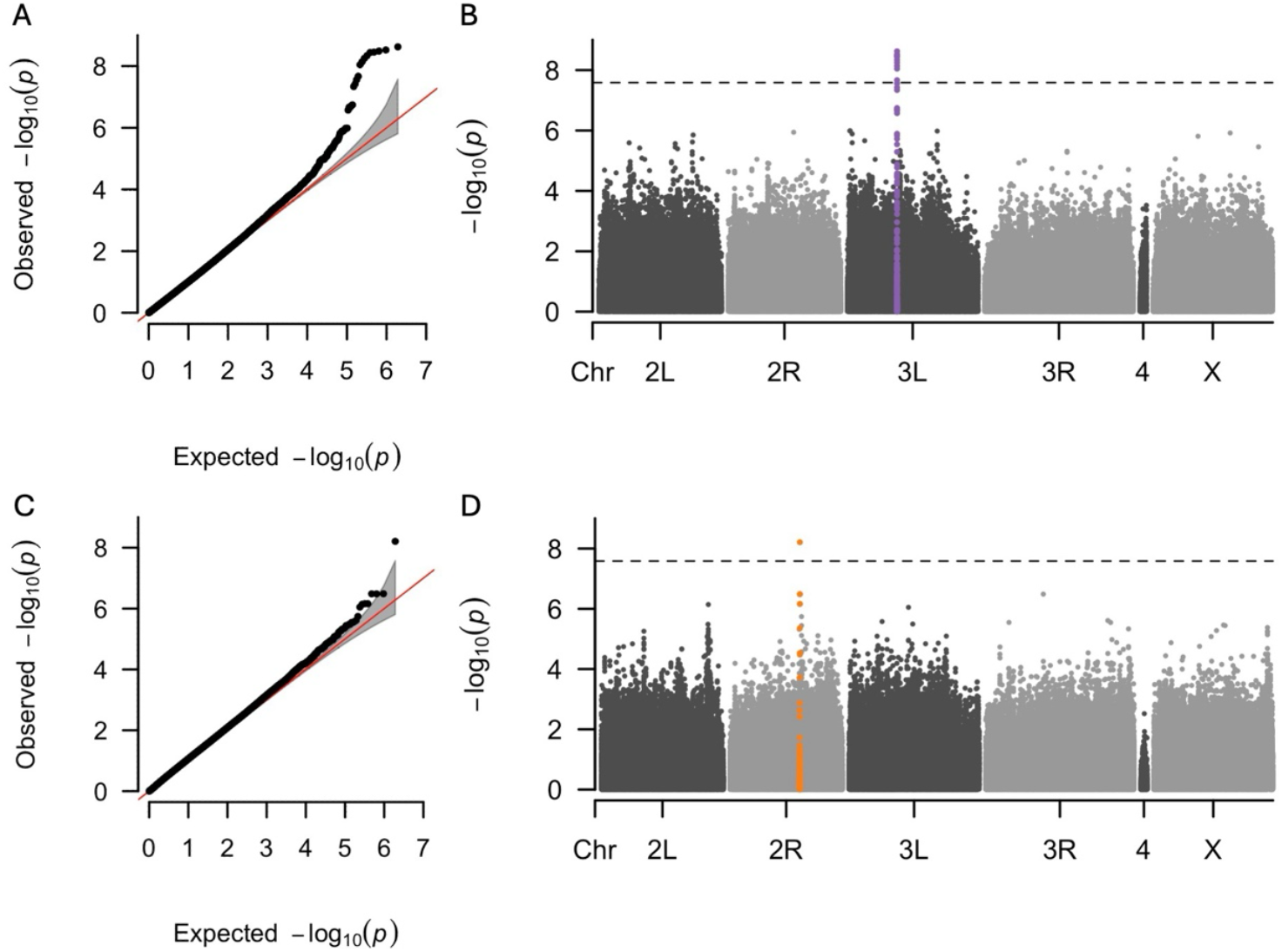
Q-Q and Manhattan plots for GWAS of GDI for dry weight and lifespan. Results of GWAS for the genotype-by-diet interaction for dry weight of 7 day-old flies (**A, B**) and genotype by diet interaction for lifespan (**C, D**). Panels (**A, C**) are Q-Q plots comparing the observed -log_10_(*p*) values of each variant to the expected values, with the red line representing the null expectation and the grey area indicating the confidence interval. Panels (**B, D**) are Manhattan plots where each point represents a variant. The *y*-axes show the strength of the association between individual variants and GDI for dry weight and lifespan, expressed as -log_10_(*p*). The dashed horizontal lines indicate the significance threshold adjusted for multiple testing using the Bonferroni correction. Variants highlighted for dry weight (purple) and lifespan (orange) indicate the GWAS index variant and all other variants within ± 2500 base pairs of the index variant.

## Discussion

Dietary restriction (DR), defined as reducing calorie intake without causing malnutrition, has been shown to delay aging in numerous species (55). While some studies have shown that DR extends longevity in flies (11,56,57), Bak et al. (8) found that increasing DR reduces both longevity and healthspan. In addition, dietary responses depend on genetic background (25,27,29). Diet plays a pivotal role in promoting healthy living, yet individual responses to dietary interventions can vary widely, complicating our understanding of its effects. The aim of the current study was to explore how individual genotypes influence responses to diet and the aging process in *D. melanogaster*. We used 98 DGRP lines to examine how a standard caloric-rich diet and a restricted diet impacted lifespan and various healthspan metrics, including locomotor activity, heat knockdown time (HKDT), and dry weight.

We found that feeding DGRP flies the restricted diet resulted in an average decrease in mean lifespan compared to flies fed a nutritionally rich control diet (Figures 3A and S2A-B, Tables 2 and S10). This suggests that the restricted diet may have caused malnutrition rather than DR. Nakagawa et al. (58) conducted a meta-analysis, including a range of species, which showed a U-shaped relationship between the risk of death and amount of DR, indicating that an intermediate level of calorie intake increases lifespan. Our findings align with the broader understanding that DR can lead to reduced longevity in some contexts, possibly due to the stress associated with insufficient nutrient intake. Furthermore, mice studies on fasting, defined as a period without calorie consumption, have shown that the positive effects of DR can, in some cases, be attributed more to the effects of fasting itself rather than just the reduction in calorie intake (59,60). When *M. musculus* were exposed to fasting in DR studies, their lifespan increased, while mice exposed to an *ad libitum* diet with nondigestible fiber did not have an increased lifespan (59,60). These observations are consistent with our results showing that flies exposed to non-fasting DR using indigestible cellulose for dilution had reduced lifespans. However, Carey et al. (43) found an increase in *D. melanogaster* lifespan with DR also using an *ad libitum* feeding and calorie restriction by diluting with indigestible cellulose. Furthermore, Lenhart et al. (61) found that the effects of fasting in *D. melanogaster* depend on genetic background and do not always confer positive effects. These contrasting findings suggest complex and genotype-specific responses to diet. The healthspan-related traits assessed in our study (dry weight, locomotor activity and HKDT) showed mixed results when exposed to different diets. When flies were exposed to a restricted diet, mean dry weight decreased (Figures 3A and S2A-B, Tables 2 and S10), locomotor activity increased (Figures 3B and S2C-D, Tables 2, S8 and S14), and the directional change in HKDT depended on the age of the flies (Figures 3C and S3E-F, Tables 2, S9 and S15). These effects of diet on healthspan metrics and lifespan show that aging is a complex and context-dependent phenomenon. Our finding that dietary nutritional conditions can have opposite, and trait- and genotype-specific impacts on lifespan and healthspan challenge our ability to make general recommendations as to which conditions are most advantageous.

Interestingly, the phenotypic correlations showed that lifespan on either diet is only weakly associated with most of the healthspan traits (Figure 4 and Tables S16-S17). This finding is inconsistent with Soo et al. (40), who reported a positive correlation between stress resistance, including heat stress, and lifespan in *C. elegans*. However, the lack of correlation between locomotor activity and lifespan aligns with Wilson et al. (11), who found no correlation between climbing ability and lifespan in the DGRP under dietary restriction. Therefore, we do not see much evidence of constraint among these traits, suggesting they can respond to selection independently. Consequently, dry weight, HKDT, and locomotor activity might not be crucial for survival under nutrient-limited laboratory conditions. Additionally, we do not find evidence of antagonistic pleiotropy, with the exception of dry weight that was found to be negatively correlated with HKDT of 16 day-old flies fed on control diet.

Our study aimed to investigate trait variation in flies under different dietary conditions, focusing on the genetic background of DGRP lines. The 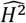 of lifespan was 0.39 on the control diet and 0.53 on the restricted diet (Figure 4 and Tables S16-S17), which are comparable to previous *D. melanogaster* studies 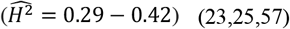. Dry weight 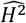 ranged from 0.25 to 0.34, aligning with the DGRP study by Jumbo-Lucioni et al. (62), but lower than Watanabe (63) who found 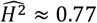. Locomotor activity 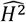 ranged from 0.37 to 0.43, consistent with other DGRP studies 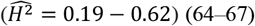. HKDT 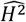 was between 0.12 and 0.36, also comparable to other DGRP findings 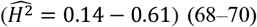. This suggests considerable genetic variation for all the traits.

Environmental factors can influence genetic variation through environment-dependent gene action. This means that genes affecting a particular trait in one environment might not be significant for that trait in another environment. This phenomenon is often reflected in altered genetic correlations, either between different traits within a single environment or the same trait across various environments (71–73). Our findings indicate that the genetic correlations for dry weight, locomotor activity, and HKDT remain consistent across ages, suggesting that the same genes influence these traits across age-classes (Figure 4 and Tables S16-S17). However, the magnitude of these genetic correlations varies with diet. Specifically, the genetic correlations for dry weight and HKDT decrease when shifting from a control diet to a restricted diet, whereas the genetic correlation for locomotor activity increases under the same dietary change. This suggests that different regulatory pathways influence dry weight, locomotor activity, and HKDT depending on the diet. The pattern of change in genetic correlations across different diets is further validated by the highly significant GDI values for all traits (Tables S7-S9). The significance of GDI is mainly attributed to changes in the rank order of lines, rather than the magnitude of between-line genetic variance for all traits. Additionally, the GAI values were also significant for all traits (Tables S7-S9). The significance of GAI is primarily due to changes in the rank order of lines, rather than the magnitude of between-line genetic variance for all traits (Figure 6 and Table S18). This indicates that trait values for dry weight, locomotor activity, HKDT, and lifespan in flies depend on their genetic background and age. These GDI and GAI findings underscore that dietary responses vary significantly not only among individuals with different genetic backgrounds, as shown previously (25,27,29), but also across different ages within those genetic backgrounds. This indicates that the effectiveness of dietary interventions depends on age, highlighting the importance of considering both genetic background and age when assessing dietary impacts on an individual. Therefore, dietary interventions should be personalized and may need to be adjusted continuously throughout an individual’s life.

We conducted GWAS to identify genetic variants associated with dietary response of lifespan, as well as age and diet response for dry weight, locomotor activity, and HKDT (Figure 7 and Tables S5-S8). We identified a variant associated with genotype-by-diet interaction of dry weight of 7 day-old flies (Figures 7A-B and Table S19), located at the intronic *PGRP-LC* locus. *PGRP-LC* encodes a transmembrane protein that acts as a pattern recognition receptor, identifying peptidoglycan in bacterial cell walls (74). It influences cellular immune responses by activating blood cells and enhancing their circulation independently of the Relish pathway, thereby playing a role in both humoral and cellular immunity (50). Activation of *PGRP-LC* in the IMD pathway in the fat body leads to the production of antimicrobial peptides during infections (51). However, the current literature does not indicate that *PGRP-LC* directly regulates fat body metabolism or functions beyond its immune response role. Furthermore, *PGRP-LC* has been found to regulate lifespan but in a temperature-dependent manner in the DGRP (25). This suggests that *PGRP-LC* is important for immune regulation rather than metabolic functions in the fat body (75). We also identified a QTL for the genotype-by-diet interaction affecting the lifespan of *D. melanogaster* (Figures 7C-D and Table S20) at the *mbl* locus. The *mbl* gene encodes an RNA-binding protein crucial for RNA metabolism, including alternative splicing and microRNA (miRNA) biogenesis (52–54). It is essential for muscle, eye, and neural tissue differentiation and plays a significant role in cardiac function, influencing lifespan in *D. melanogaster* (52,76). Overexpression of *mbl* in myotonic dystrophy type 1 (*DM1*) models improved muscle function and lifespan (77). Manipulating expression of *mbl* and its counterpart *Bru-3* led to asynchronous heartbeats and dilated cardiomyopathy, highlighting the role of *mbl* in cardiac health (78,79). These findings suggest that the regulatory functions of *mlb* in RNA processing may impact lifespan and aging in a diet-dependent manner. The candidate genes have only been identified in genotype-by-diet interactions, highlighting the crucial role of genetics in shaping the aging response to diet. The GWAS results presented here indicate two candidate genes, *PGRP-LC* and *mbl*, which can be targeted for further research on aging.

## Author Contributions

Conceptualization, N.K.B., T.N.K. and P.D.R.; Data Curation, N.K.B., Formal Analysis, N.K.B., T.F.C.M., F.M. and P.D.R.; Funding Acquisition, T.N.K., P.D.R., T.F.C.M. and F.M.; Investigation, N.K.B. and T.N.K; Methodology, N.K.B., T.N.K. and P.D.R.; Project Administration, N.K.B. and T.N.K.; Resources, T.N.K., P.D.R. and T.F.C.M.; Supervision, T.N.K., P.D.R., T.F.C.M. and F.M.; Visualization, N.K.B.; Writing – Original Draft Preparation, N.K.B., T.N.K. and P.D.R.; Writing – Review and Editing, T.F.C.M., F.M.; K.L.N. and J.L.N.

## Funding

This research was funded by the Danish Council for Independent Research (DFF-2032-00205A) to T.N.K. and P.D.R, and by the United States National Institutes of Health under award numbers P20 GM139769 and R01 AG073181 to T.F.C.M., and R35GM146868 to F.M. The content is solely the responsibility of the authors and does not necessarily represent the official views of the Danish Council for Independent Research and the National Institutes of Health. The funders had no role in the decision to publish or prepare the manuscript.

## Acknowledgements

We thank Andreas Mølgaard Andersen, Anna Klausholt Bak, Christian Dupont Danielsen, Emilie Battersby, Frederik Kjær, Helle Blendstrup, Jane Pries Jakobsen, Jonas Bruhn Wesseltoft, Julie Wachmann Jensen, Madeleine Picard Svejgaard Madsen, Magnus Tudsborg Frantzen, Michael Ørsted, Selma Andersen, Stine Frey Laursen, Susan Marie Hansen and Timo Kirwa for technical assistance in the laboratory. With thanks to Vijay Shankar for assistance with GWAS analysis.

## Conflicts of interest

The authors declare no conflicts of interest.

## Notes

### Competing Interest Statement

The authors have declared no competing interest.

## References

1. Raymond Pearl. Experiments on Longevity. Q Rev Biol. 1928 May 9;3(3):391– 407.

2. Percy Baumberger J. Studies in the Longevity of Insects. Ann Entomol Soc Am. 1914 Dec 1;7(4):323–53.

3. George LK, Palmore E, Cohen HJ. The Duke Center for the Study of Aging: One of Our Earliest Roots. Gerontologist. 2014 Feb 1;54(1):59–66.

4. Le Bourg E. Lifespan Versus Healthspan. In 2020. p. 439–52.

5. Seals DR, Justice JN, LaRocca TJ. Physiological geroscience: targeting function to increase healthspan and achieve optimal longevity. J Physiol. 2016 Apr 15;594(8):2001–24.

6. Kirkland JL, Peterson C. Healthspan, Translation, and New Outcomes for Animal Studies of Aging. J Gerontol A Biol Sci Med Sci. 2009 Feb 1;64A(2):209–12.

7. Tissenbaum HA. Genetics, Life Span, Health Span, and the Aging Process in Caenorhabditis elegans. J Gerontol A Biol Sci Med Sci. 2012 May 1;67A(5):503–10.

8. Bak NK, Rohde PD, Kristensen TN. Strong Sex-Dependent Effects of Malnutrition on Life- and Healthspan in Drosophila melanogaster. Insects. 2023 Dec 26;15(1):9.

9. Wu RT, Cao L, Mattson E, Witwer KW, Cao J, Zeng H, et al. Opposing impacts on healthspan and longevity by limiting dietary selenium in telomere dysfunctional mice. Aging Cell. 2017 Feb 21;16(1):125–35.

10. Fischer KE, Hoffman JM, Sloane LB, Gelfond JAL, Soto VY, Richardson AG, et al. A cross-sectional study of male and female C57BL/6Nia mice suggests lifespan and healthspan are not necessarily correlated. Aging. 2016 Oct 2;8(10):2370–91.

11. Wilson KA, Beck JN, Nelson CS, Hilsabeck TA, Promislow D, Brem RB, et al. GWAS for Lifespan and Decline in Climbing Ability in Flies upon Dietary Restriction Reveal decima as a Mediator of Insulin-like Peptide Production. Current Biology. 2020 Jul;30(14):2749-2760.e3.

12. World Bank Group. https://data.worldbank.org/indicator/SP.DYN.LE00.IN? 2022. Life expectancy at birth, total (years).

13. Cambois E, Blachier A, Robine JM. Aging and health in France: an unexpected expansion of disability in mid-adulthood over recent years. The European Journal of Public Health. 2013 Aug 1;23(4):575–81.

14. Jagger C, Gillies C, Moscone F, Cambois E, Van Oyen H, Nusselder W, et al. Inequalities in healthy life years in the 25 countries of the European Union in 2005: a cross-national meta-regression analysis. The Lancet. 2008 Dec;372(9656):2124–31.

15. Garmany A, Yamada S, Terzic A. Longevity leap: mind the healthspan gap. NPJ Regen Med. 2021 Sep 23;6(1):57.

16. Hoffman JM, Valencak TG. A short life on the farm: aging and longevity in agricultural, large-bodied mammals. Geroscience. 2020 Jun 2;42(3):909–22.

17. Comizzoli P, Ottinger MA. Understanding Reproductive Aging in Wildlife to Improve Animal Conservation and Human Reproductive Health. Front Cell Dev Biol. 2021 May 19;9.

18. Ottinger MA, Grace JK, Maness TJ. Global challenges in aging: insights from comparative biology and one health. Frontiers in Toxicology. 2024 May 30;6.

19. Kirkwood TBL. Understanding the Odd Science of Aging. Cell. 2005 Feb;120(4):437–47.

20. Flatt T, Partridge L. Horizons in the evolution of aging. BMC Biol. 2018 Dec 20;16(1):93.

21. Krittika S, Yadav P. Dietary protein restriction deciphers new relationships between lifespan, fecundity and activity levels in fruit flies Drosophila melanogaster. Sci Rep. 2020 Jun 22;10(1):10019.

22. Matta ME, Tribuzio CA, Davidson LNK, Fuller KR, Dunne GC, Andrews AH. A review of the Pacific sleeper shark Somniosus pacificus: biology and fishery interactions. Polar Biol. 2024 May 14;47(5):433–58.

23. Ivanov DK, Escott-Price V, Ziehm M, Magwire MM, Mackay TFC, Partridge L, et al. Longevity GWAS Using the Drosophila Genetic Reference Panel. J Gerontol A Biol Sci Med Sci. 2015 Dec;70(12):1470–8.

24. Urban ND, Cavataio JP, Berry Y, Vang B, Maddali A, Sukpraphrute RJ, et al. Explaining inter-lab variance in C. elegans N2 lifespan: Making a case for standardized reporting to enhance reproducibility. Exp Gerontol. 2021 Dec;156:111622.

25. Huang W, Campbell T, Carbone MA, Jones WE, Unselt D, Anholt RRH, et al. Context-dependent genetic architecture of Drosophila life span. PLoS Biol. 2020 Mar 5;18(3):e3000645.

26. Mackay TFC, Huang W. Charting the genotype-phenotype map: lessons from the Drosophila melanogaster Genetic Reference Panel. Wiley Interdiscip Rev Dev Biol. 2018 Jan;7(1).

27. Francis D, Ghazanfar S, Havula E, Krycer JR, Strbenac D, Senior A, et al. Genome-wide analysis in Drosophila reveals diet-by-gene interactions and uncovers diet-responsive genes. G3: Genes, Genomes, Genetics. 2021 Oct 1;11(10).

28. Duun Rohde P, Krag K, Loeschcke V, Overgaard J, Sørensen P, Nygaard Kristensen T. A Quantitative Genomic Approach for Analysis of Fitness and Stress Related Traits in a Drosophila melanogaster Model Population. Int J Genomics. 2016;2016.

29. Patel SP, Talbert ME. Identification of genetic modifiers of lifespan on a high sugar diet in the Drosophila Genetic Reference Panel. Heliyon. 2021 Jun 1;7(6).

30. Unckless RL, Rottschaefer SM, Lazzaro BP. The Complex Contributions of Genetics and Nutrition to Immunity in Drosophila melanogaster. PLoS Genet. 2015 Mar 12;11(3).

31. Mackay TFC, Anholt RRH. Pleiotropy, epistasis and the genetic architecture of quantitative traits. Vol. 25, Nature Reviews Genetics. Nature Research; 2024. p. 639–57.

32. Jin K, Wilson KA, Beck JN, Nelson CS, Brownridge GW, Harrison BR, et al. Correction: Genetic and metabolomic architecture of variation in diet restriction-mediated lifespan extension in Drosophila. Vol. 18, PLoS Genetics. Public Library of Science; 2022.

33. Jordan KW, Morgan TJ, Mackay TFC. Quantitative trait loci for locomotor behavior in Drosophila melanogaster. Genetics. 2006;174(1):271–84.

34. Krittika S, Yadav P. Trans-generational effect of protein restricted diet on adult body and wing size of Drosophila melanogaster. R Soc Open Sci. 2022;9(1).

35. Eickelberg V, Lüersen K, Staats S, Rimbach G. Phenotyping of Drosophila melanogaster—A Nutritional Perspective. Vol. 12, Biomolecules. MDPI; 2022.

36. Bubliy OA, Loeschcke V. Correlated responses to selection for stress resistance and longevity in a laboratory population of Drosophila melanogaster. In: Journal of Evolutionary Biology. 2005. p. 789–803.

37. Belyi AA, Alekseev AA, Fedintsev AY, Balybin SN, Proshkina EN, Shaposhnikov M V., et al. The Resistance of Drosophila melanogaster to Oxidative, Genotoxic, Proteotoxic, Osmotic Stress, Infection, and Starvation Depends on Age According to the Stress Factor. Antioxidants. 2020 Dec 7;9(12):1239.

38. Banse SA, Jackson EG, Sedore CA, Onken B, Hall D, Coleman-Hulbert A, et al. The coupling between healthspan and lifespan in Caenorhabditis depends on complex interactions between compound intervention and genetic background. Aging. 2024 Apr 12;

39. Badial K, Lacayo P, Murakami S. Biology of healthy aging: Biological hallmarks of stress resistance-related and unrelated to longevity in humans [Internet]. 2024. Available from: http://biorxiv.org/lookup/doi/10.1101/2024.07.19.604380

40. Soo SK, Traa A, Rudich ZD, Moldakozhayev A, Mistry M, Van Raamsdonk JM. Genetic basis of enhanced stress resistance in long-lived mutants highlights key role of innate immunity in determining longevity. Aging Cell. 2023 Feb 1;22(2).

41. MacKay TFC, Richards S, Stone EA, Barbadilla A, Ayroles JF, Zhu D, et al. The Drosophila melanogaster Genetic Reference Panel. Nature. 2012 Feb 9;482(7384):173–8.

42. Huang W, Massouras A, Inoue Y, Peiffer J, Ràmia M, Tarone AM, et al. Natural variation in genome architecture among 205 Drosophila melanogaster Genetic Reference Panel lines. Genome Res. 2014;24(7):1193–208.

43. Carey MR, Archer CR, Rapkin J, Castledine M, Jensen K, House CM, et al. Mapping sex differences in the effects of protein and carbohydrates on lifespan and reproduction in Drosophila melanogaster: is measuring nutrient intake essential? Biogerontology. 2022 Feb 1;23(1):129–44.

44. Strilbytska O, Yurkevych I, Semaniuk U, Gospodaryov D, Simpson SJ, Lushchak O. Life-History Trade-Offs in Drosophila : Flies Select a Diet to Maximize Reproduction at the Expense of Lifespan. J Gerontol A Biol Sci Med Sci. 2024 May 1;79(5).

45. Jensen K, McClure C, Priest NK, Hunt J. Sex-specific effects of protein and carbohydrate intake on reproduction but not lifespan in Drosophila melanogaster. Aging Cell. 2015 Aug;14(4):605–15.

46. Rohde PD, Fourie Sørensen I, Sørensen P. Qgg: An R package for large-scale quantitative genetic analyses. Bioinformatics. 2020 Apr 15;36(8):2614–5.

47. Mackay T, Falconer D. Introduction to Quantitative Genetics. 4th ed. Pearson Education Limited; 1996.

48. Cockerham C. Statistical Genetics and Plant Breeding. Hanson W, Robinson R, editors. Washington, D.C.: National Academies Press; 1963. 53–94 p.

49. Gnambs T. A Brief Note on the Standard Error of the Pearson Correlation. Collabra Psychol. 2023 Sep 6;9(1).

50. Borge-Renberg K. Communicate or die: Signaling in Drosophila immunity. 1st ed. Vol. 1. Umeå centrum för molekylär patogenes (UCMP) (Medicinska fakulteten); 2008. 1–63 p.

51. Aggarwal K, Silverman N. Positive and negative regulation of the Drosophila immune response. BMB Rep. 2008 Apr 30;41(4):267–77.

52. Li JSS, Millard SS. Deterministic splicing of Dscam2 is regulated by Muscleblind. Sci Adv. 2019 Jan 4;5(1).

53. Oddo JC, Saxena T, McConnell OL, Berglund JA, Wang ET. Conservation of context-dependent splicing activity in distant Muscleblind homologs. Nucleic Acids Res. 2016 Sep 30;44(17):8352–62.

54. Fernandez-Costa JM, Llamusi MB, Garcia-Lopez A, Artero R. Alternative splicing regulation by Muscleblind proteins: from development to disease. Biological Reviews. 2011 Nov;86(4):947–58.

55. Le Couteur DG, Raubenheimer D, Solon-Biet S, de Cabo R, Simpson SJ. Does diet influence aging? Evidence from animal studies. J Intern Med. 2024 Apr 24;295(4):400–15.

56. Li M, Macro J, Huggins BJ, Meadows K, Mishra D, Martin D, et al. Extended lifespan in female Drosophila melanogaster through late-life calorie restriction. Geroscience. 2024 Jul 2;46(5):4017–35.

57. Durham MF, Magwire MM, Stone EA, Leips J. Genome-wide analysis in Drosophila reveals age-specific effects of SNPs on fitness traits. Nat Commun. 2014 Jul 8;5(1):4338.

58. Nakagawa S, Lagisz M, Hector KL, Spencer HG. Comparative and meta-analytic insights into life extension via dietary restriction. Aging Cell. 2012 Jun 22;11(3):401–9.

59. Pak HH, Haws SA, Green CL, Koller M, Lavarias MT, Richardson NE, et al. Fasting drives the metabolic, molecular and geroprotective effects of a calorie-restricted diet in mice. Nat Metab. 2021 Oct 18;3(10):1327–41.

60. Solon-Biet SM, McMahon AC, Ballard JWO, Ruohonen K, Wu LE, Cogger VC, et al. The Ratio of Macronutrients, Not Caloric Intake, Dictates Cardiometabolic Health, Aging, and Longevity in Ad Libitum-Fed Mice. Cell Metab. 2014 Mar;19(3):418–30.

61. Lenhart BA, Ahsan A, McHaty M, Bergland AO. Improvement of starvation resistance via periodic fasting is genetically variable in Drosophila melanogaster. Physiol Entomol. 2024 Sep 4;49(3):270–8.

62. Jumbo-Lucioni P, Ayroles JF, Chambers MM, Jordan KW, Leips J, Mackay TF, et al. Systems genetics analysis of body weight and energy metabolism traits in Drosophila melanogaster. BMC Genomics. 2010 Dec 11;11(1):297.

63. Watanabe LP, Riddle NC. GWAS reveal a role for the central nervous system in regulating weight and weight change in response to exercise. Sci Rep. 2021 Mar 4;11(1):5144.

64. Videlier M, Rundle HD, Careau V. Sex-specific genetic (co)variances of standard metabolic rate, body mass and locomotor activity in Drosophila melanogaster. J Evol Biol. 2021 Aug 24;34(8):1279–89.

65. Jordan KW, Carbone MA, Yamamoto A, Morgan TJ, Mackay TF. Quantitative genomics of locomotor behavior in Drosophila melanogaster. Genome Biol. 2007 Aug 21;8(8):R172.

66. Rohde PD, Østergaard S, Kristensen TN, Sørensen P, Loeschcke V, Mackay TFC, et al. Functional Validation of Candidate Genes Detected by Genomic Feature Models. G3 Genes|Genomes|Genetics. 2018 May 1;8(5):1659–68.

67. Noer NK, Rohde PD, Sørensen P, Bahrndorff S, Kristensen TN. Diurnal variation in genetic parameters for locomotor activity in Drosophila melanogaster assessed under natural thermal conditions. J Evol Biol. 2024 Mar 1;37(3):336–45.

68. Rolandi C, Lighton JRB, de la Vega GJ, Schilman PE, Mensch J. Genetic variation for tolerance to high temperatures in a population of Drosophila melanogaster. Ecol Evol. 2018 Nov 11;8(21):10374–83.

69. Lecheta MC, Awde DN, O’Leary TS, Unfried LN, Jacobs NA, Whitlock MH, et al. Integrating GWAS and Transcriptomics to Identify the Molecular Underpinnings of Thermal Stress Responses in Drosophila melanogaster. Front Genet. 2020 Jun 23;11.

70. Soto J, Pinilla F, Olguín P, Castañeda LE. Genetic architecture of the thermal tolerance landscape of Drosophila melanogaster. 2024.

71. Sgrò CM, Hoffmann AA. Genetic correlations, tradeoffs and environmental variation. Heredity (Edinb). 2004 Sep 28;93(3):241–8.

72. Vieira C, Pasyukova EG, Zeng ZB, Hackett JB, Lyman RF, Mackay TFC. Genotype-Environment Interaction for Quantitative Trait Loci Affecting Life Span in Drosophila melanogaster. Genetics. 2000 Jan 1;154(1):213–27.

73. Agrawal AF, Stinchcombe JR. How much do genetic covariances alter the rate of adaptation? Proceedings of the Royal Society B: Biological Sciences. 2009 Mar 22;276(1659):1183–91.

74. Kaneko T, Yano T, Aggarwal K, Lim JH, Ueda K, Oshima Y, et al. PGRP-LC and PGRP-LE have essential yet distinct functions in the Drosophila immune response to monomeric DAP-type peptidoglycan. Nat Immunol. 2006 Jul 11;7(7):715–23.

75. Gendrin M, Zaidman-Rémy A, Broderick NA, Paredes J, Poidevin M, Roussel A, et al. Functional Analysis of PGRP-LA in Drosophila Immunity. PLoS One. 2013 Jul 26;8(7):e69742.

76. Mutsuddi M, Marshall CM, Benzow KA, Koob MD, Rebay I. The Spinocerebellar Ataxia 8 Noncoding RNA Causes Neurodegeneration and Associates with Staufen in Drosophila. Current Biology. 2004 Feb;14(4):302–8.

77. Cerro-Herreros E, Fernandez-Costa JM, Sabater-Arcis M, Llamusi B, Artero R. Derepressing muscleblind expression by miRNA sponges ameliorates myotonic dystrophy-like phenotypes in Drosophila. Sci Rep. 2016 Nov 2;6(1):36230.

78. Souidi A, Nakamori M, Zmojdzian M, Jagla T, Renaud Y, Jagla K. Deregulations of miR-1 and its target Multiplexin promote dilated cardiomyopathy associated with myotonic dystrophy type 1. EMBO Rep. 2023 Apr 5;24(4).

79. Auxerre-Plantié E, Nakamori M, Renaud Y, Huguet A, Choquet C, Dondi C, et al. Straightjacket/α2δ3 deregulation is associated with cardiac conduction defects in myotonic dystrophy type 1. Elife. 2019 Dec 12;8.

