## Supplementary Material 1 for "The Role of Genetic Variation in Shaping Phenotypic Responses to Diet in Aging *Drosophila melanogaster*"

[¶] These authors contributed equally

---

This supplementary file contains:

**Supplementary Figures**

| | |
|---|---|
| Figure S1 | Correlations of estimated heritability between all data and balanced data |
| Figure S2 | Histogram plots |
| Figure S3 | Survival analysis of lifespan |
| Figure S4 | Reaction norm plots across ages |
| Figure S5 | Q-Q plot and Manhattan plot for dry weight |
| Figure S6 | Q-Q plot and Manhattan plot for locomotor activity |
| Figure S7 | Q-Q plot and Manhattan plot for HKDT |
| Figure S8 | Q-Q plot and Manhattan plot for lifespan |

**Supplementary Tables**

| | |
|---|---|
| Table S1 | Number of flies and lines on balanced and unbalanced data of dry weight |
| Table S2 | Number of flies and lines on balanced and unbalanced data of locomotor activity |
| Table S3 | Number of flies and lines on balanced and unbalanced data of HKDT |
| Table S4 | Summary of ANOVA on data of dry weight |
| Table S5 | Summary of ANOVA on data of locomotor activity |
| Table S6 | Summary of ANOVA on data of HKDT |
| Table S7 | Summary of ANOVA on balanced data of dry weight |
| Table S8 | Summary of ANOVA on balanced data of locomotor activity |
| Table S9 | Summary of ANOVA on balanced data of HKDT |
| Table S10 | Line mean and standard error of dry weight for separate days and diets [**Excel**] |
| Table S11 | Summary of ANOVA on data of lifespan |
| Table S12 | Summary of Coxph on data of lifespan |
| Table S13 | Line mean and standard error of lifespan for separate days and diets [**Excel**] |
| Table S14 | Line mean and standard error of locomotor activity for separate days and diets [**Excel**] |
| Table S15 | Line mean and standard error of HKDT for separate days and diets [**Excel**] |
| Table S16 | Phenotypic correlation, heritability and genotypic correlation matrix of all traits with standard errors [**Excel**] |
| Table S17 | Phenotypic correlation, heritability and genotypic correlation matrix of all traits with 95% Confidence intervals [**Excel**] |
| Table S18 | Matrix of the percent contribution of changes in rank order of all traits [**Excel**] |
| Table S19 | GWAS results for the genotype-by-diet interaction for dry weight of 7 day-old flies [**Excel**] |
| Table S20 | GWAS results for the genotype-by-diet interaction for lifespan of flies [**Excel**] |

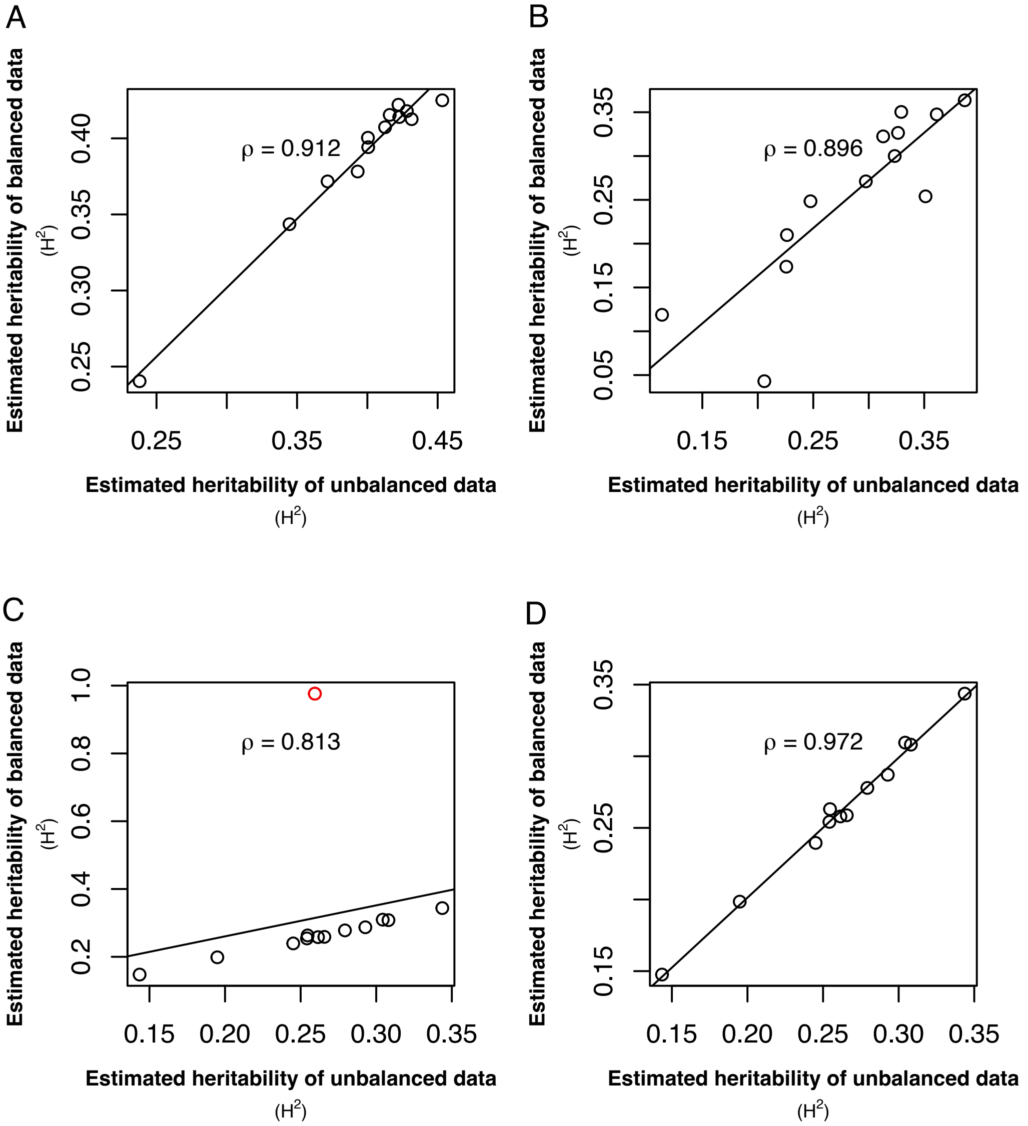


**Figure S1. Correlations of estimated heritability between all data and unbalanced data.** The balanced dataset was generated by excluding DGRP lines that were not present in both control and restricted diets, whereas the unbalanced dataset included all lines. The figures represent healthspan metrics: (**A**) locomotor activity, (**B**) heat knockdown time (HKDT), (**C**) dry weight with an outlier indicated as a red point, and (**D**) dry weight excluding the outlier. The symbol 'ρ' denotes the Spearman correlation coefficient.

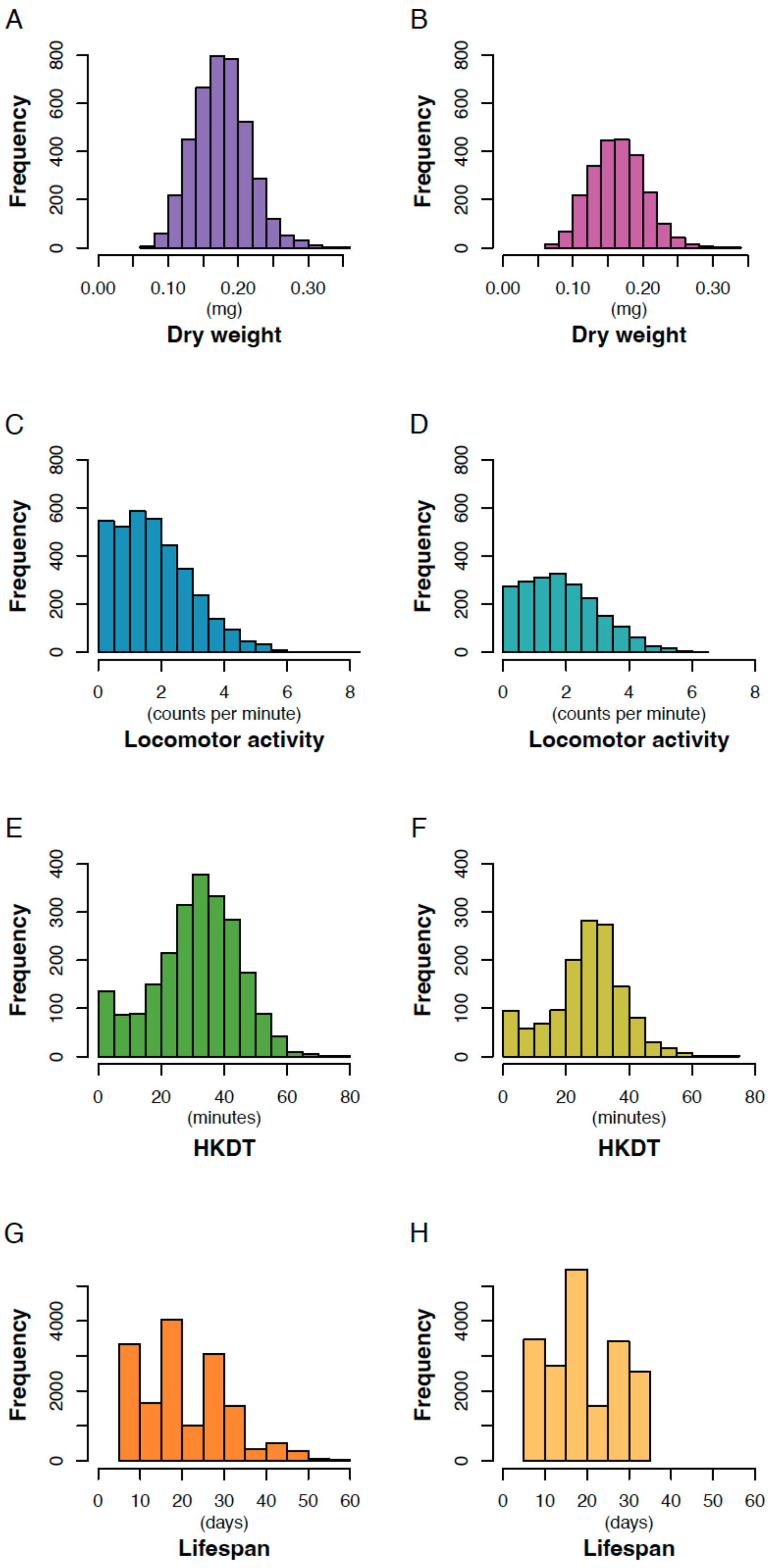


**Figure S2. Histogram plots.** Histograms for (**A**,**B**) dry weight (purple), (**C**,**D**) locomotor activity (blue), (**E**,**F**) HKDT (green) and (**G**,**H**) lifespan (orange) on control (dark shades) and restricted (light shades) diets.

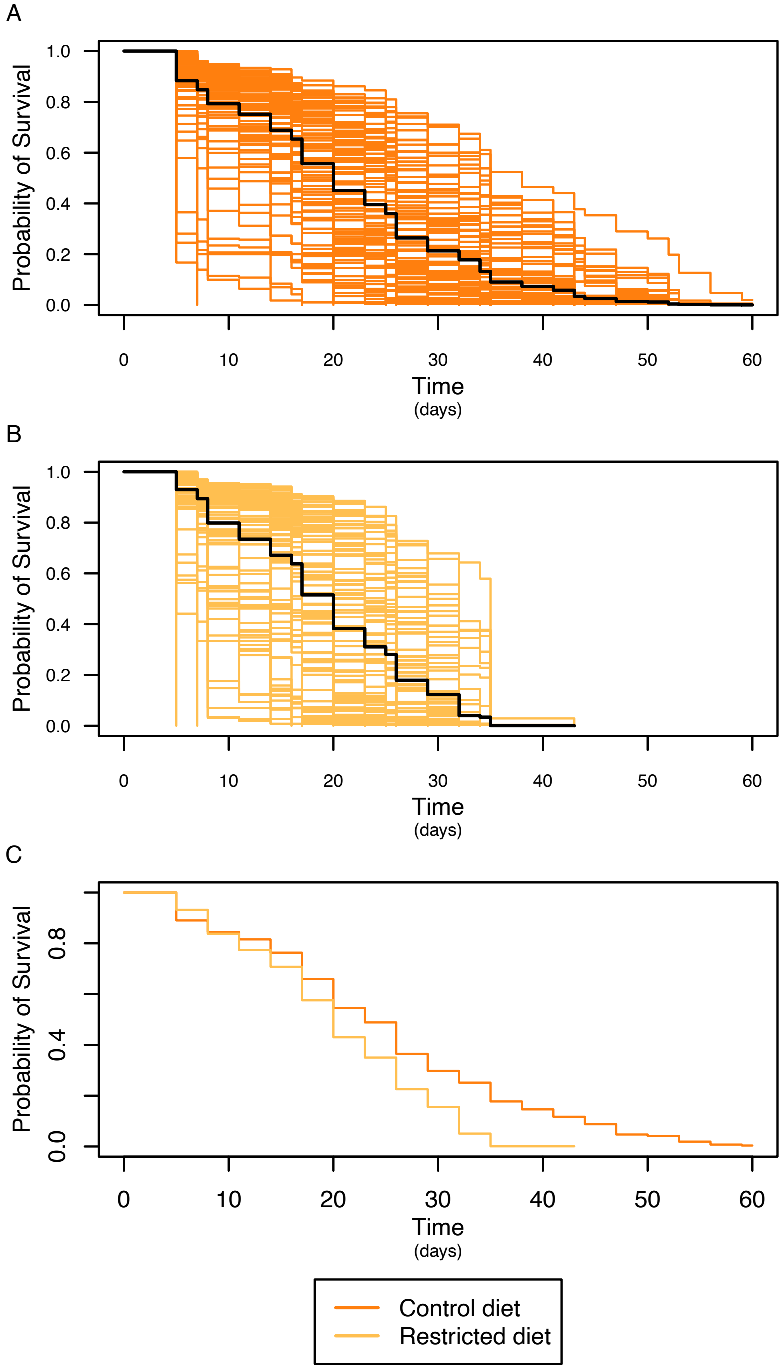


**Figure S3. Survival analysis of lifespan.** Lines fed on (**A**) control diet (dark orange), (**B**) restricted diet (light orange), where black lines indicate mean lifespan for separate diets. (**C**) mean across lines fed control diet (dark orange) and restricted diet (light orange).

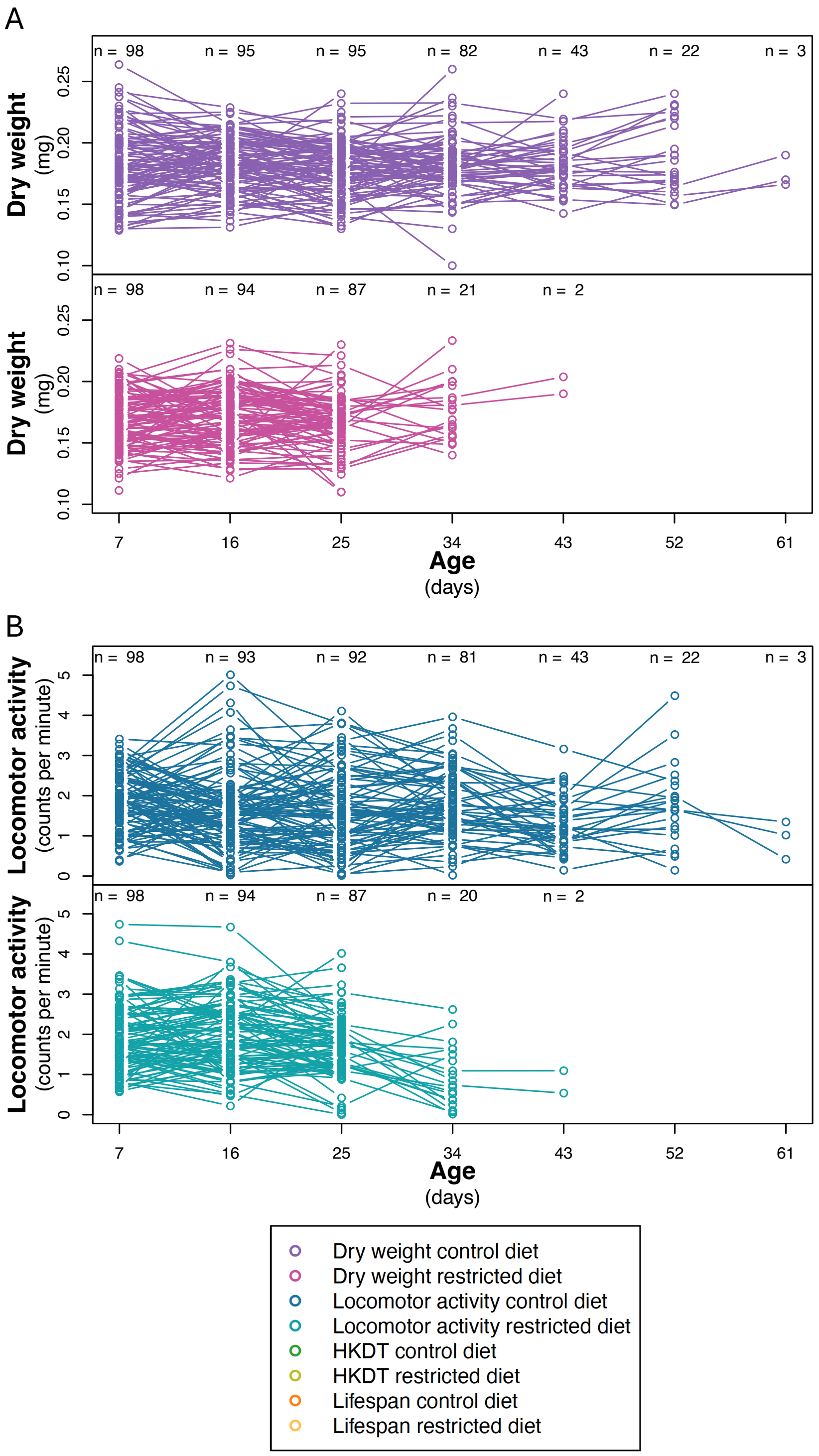


**Figure S4. Reaction norm plots across ages.** Line means of (**A**) dry weight (purple), (**B**) locomotor activity (blue) and (**C**) HKDT (green) across seven ages, fed on control (dark shades) and restricted (light shades) diets. (**D**) lifespan (orange) also fed on control (dark shades) and restricted (light shades) diets. Points represent the mean phenotype of each DGRP line, and lines connecting each points across age or diet represent the same DGRP line. Numbers (n) of DGRP lines assessed at each age and diet are indicated above the points.

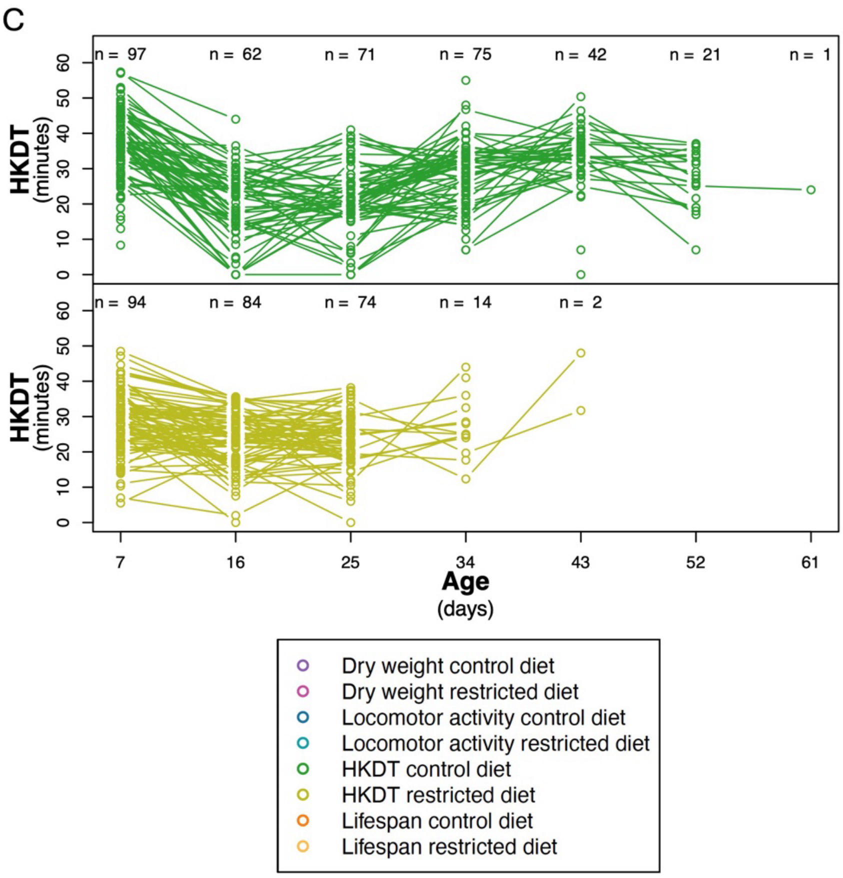


**Figure S4. Reaction norm plots across ages (continued).** Line means of (**A**) dry weight (purple), (**B**) locomotor activity (blue) and (**C**) HKDT (green) across seven ages, fed on control diet (dark shades) and restricted diet (light shades). Points represent the mean phenotype of each DGRP line, and lines connecting points across age represent the same DGRP line. Numbers (n) of DGRP lines assessed at each age and diet are indicated above the points.

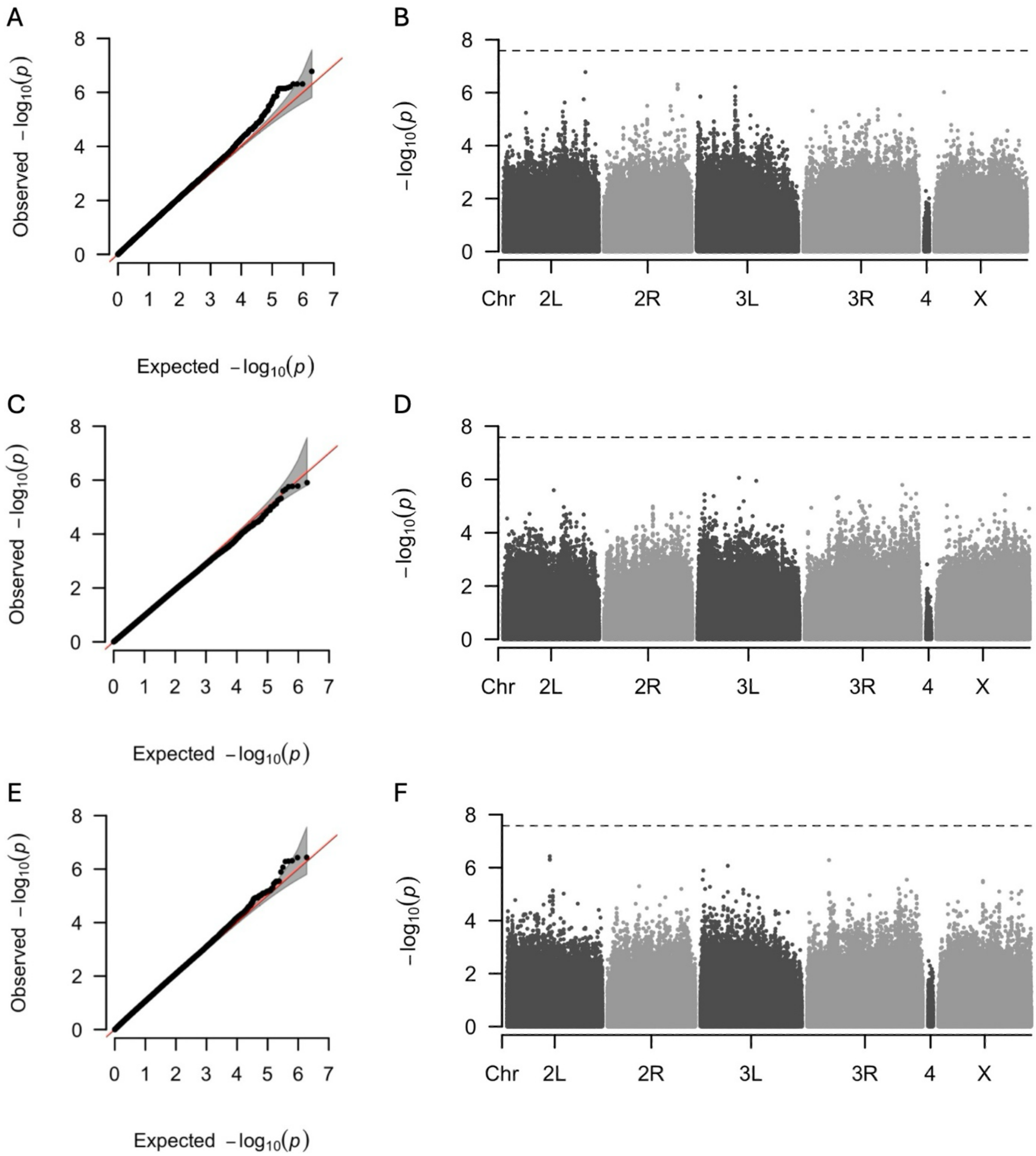


**Figure S5**. **Q-Q plot and Manhattan plot for dry weight.** Results of a GWAS to identify variants associated with dry weight of (**A**, **B**) 7 day-old flies fed the control diet, (**C**, **D**) 7 day-old flies fed the restricted diet, (**E**, **F**) 16 day-old flies fed the control diet, (**G**, **H**) 16 day-old flies fed the restricted diet, (**I**, **J**) genotype-by-diet interaction for 16 day-old flies, and (**K**, **L**) genotype-by-age interaction for flies fed the restricted diet. Panels (**A**, **C**, **E**, **G**, **I**, **K**) are Q-Q plots comparing the observed -log(*p*) values of each variant to the expected values, with the red line representing the null expectation and the grey area indicating the confidence interval. Panels (**B**, **D**, **F**, **H**, **J**, **L**) are Manhattan plots where each point represents a variant. The *y*-axis shows the strength of the association between individual variants and dry weight, expressed as -log(*p*). The dashed horizontal line indicates the significance threshold adjusted for multiple testing using the Bonferroni correction.

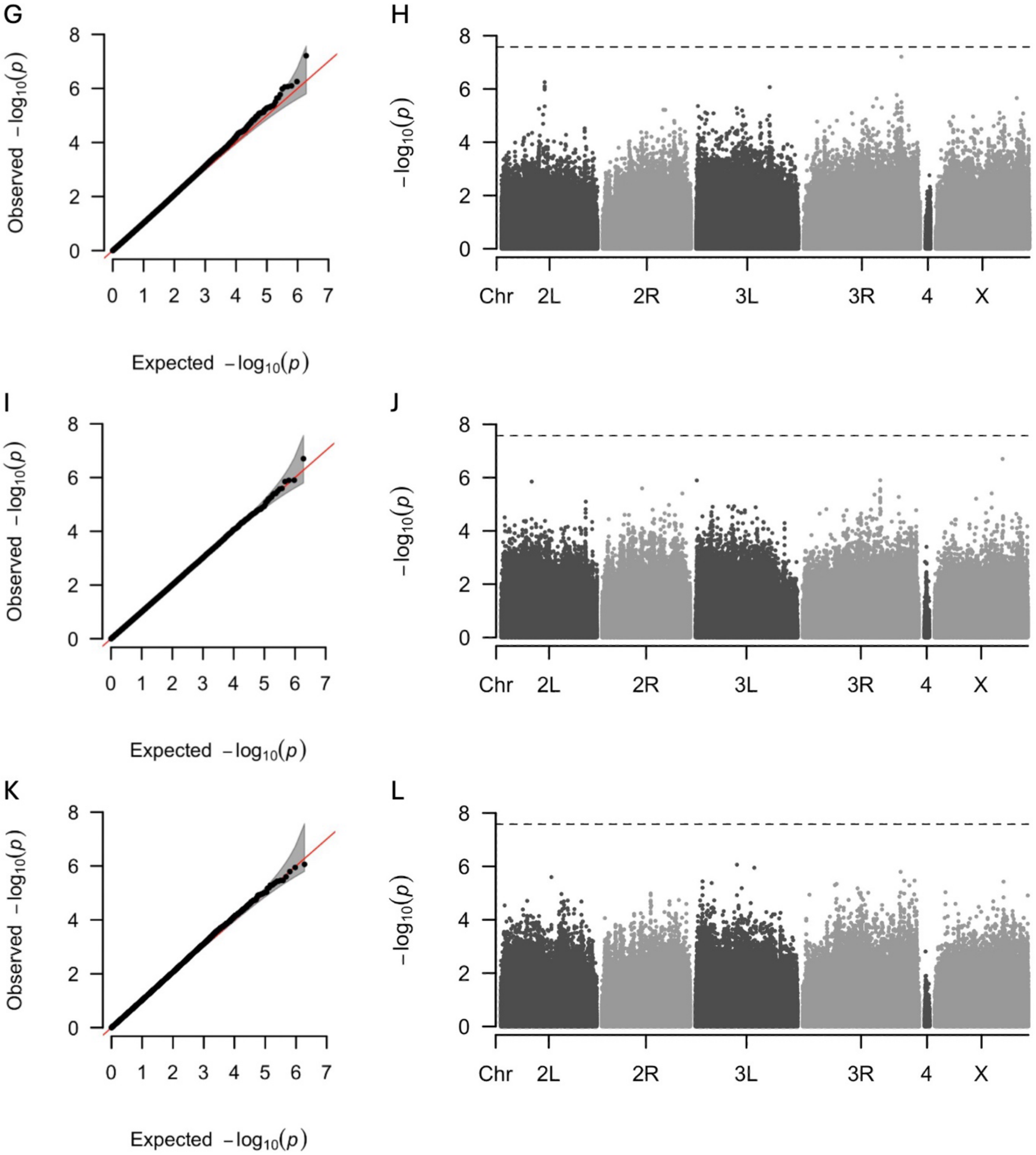


**Figure S5 (continued)**. **Q-Q plot and Manhattan plot for dry weight.** Results of a GWAS to identify variants associated with dry weight of (**A**, **B**) 7 day-old flies fed the control diet, (**C**, **D**) 7 day-old flies fed the restricted diet, (**E**, **F**) 16 day-old flies fed the control diet, (**G**, **H**) 16 day-old flies fed the restricted diet, (**I**, **J**) genotype-by-diet interaction for 16 day-old flies, and (**K**, **L**) genotype-by-age interaction for flies fed the restricted diet. Panels (**A**, **C**, **E**, **G**, **I**, **K**) are Q-Q plots comparing the observed -log($p$) values of each variant to the expected values, with the red line representing the null expectation and the grey area indicating the confidence interval. Panels (**B**, **D**, **F**, **H**, **J**, **L**) are Manhattan plots where each point represents a variant. The $y$-axis shows the strength of the association between individual variants and dry weight, expressed as -log($p$). The dashed horizontal line indicates the significance threshold adjusted for multiple testing using the Bonferroni correction.

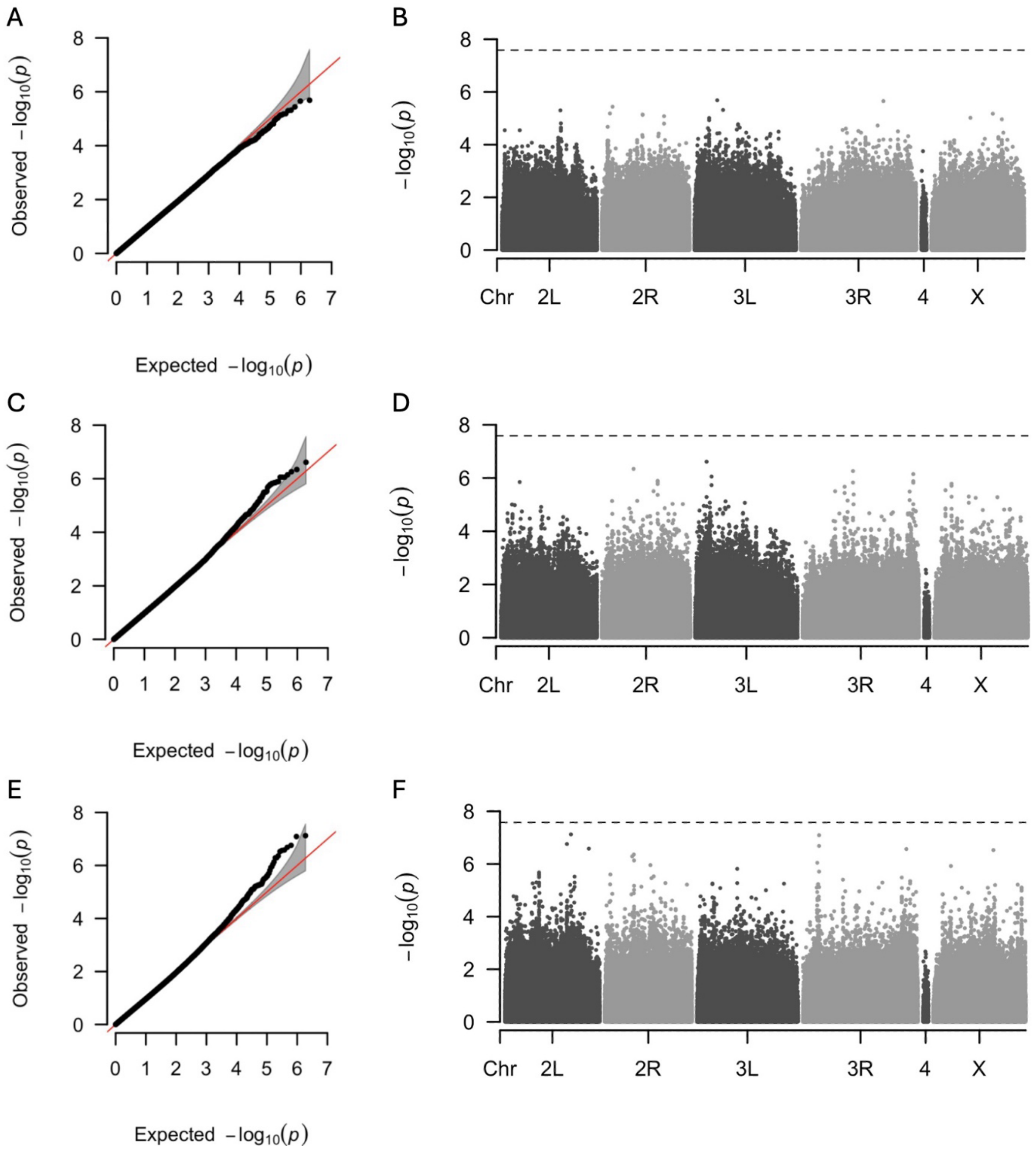


**Figure S6**. **Q-Q plot and Manhattan plot for locomotor activity.** Results of a GWAS to identify variants associated with locomotor activity of (**A**, **B**) 7 day-old flies fed the control diet, (**C**, **D**) 7 day-old flies fed the restricted diet, (**E**, **F**) 16 day-old flies fed the control diet, (**G**, **H**) 16 day-old flies fed the restricted diet, (**I**, **J**) genotype-by-diet interaction for 7 day-old flies, (**K**, **L**) genotype-by-diet interaction for 16 day-old flies, (**M**, **N**) genotype-by-age interaction for flies fed the control diet, and (**O**, **P**) genotype-by-age interaction for flies fed the restricted diet. Panels (**A**, **C**, **E**, **G**, **I**, **K, M, O**) are Q-Q plots comparing the observed -log(p) values of each variant to the expected values, with the red line representing the null expectation and the grey area indicating the confidence interval. Panels (**B**, **D**, **F**, **H**, **J**, **L**, **N**, **P**) are Manhattan plots where each point represents a variant. The y-axis shows the strength of the association between individual variants and dry weight, expressed as -log(p). The dashed horizontal line indicates the significance threshold adjusted for multiple testing using the Bonferroni correction.

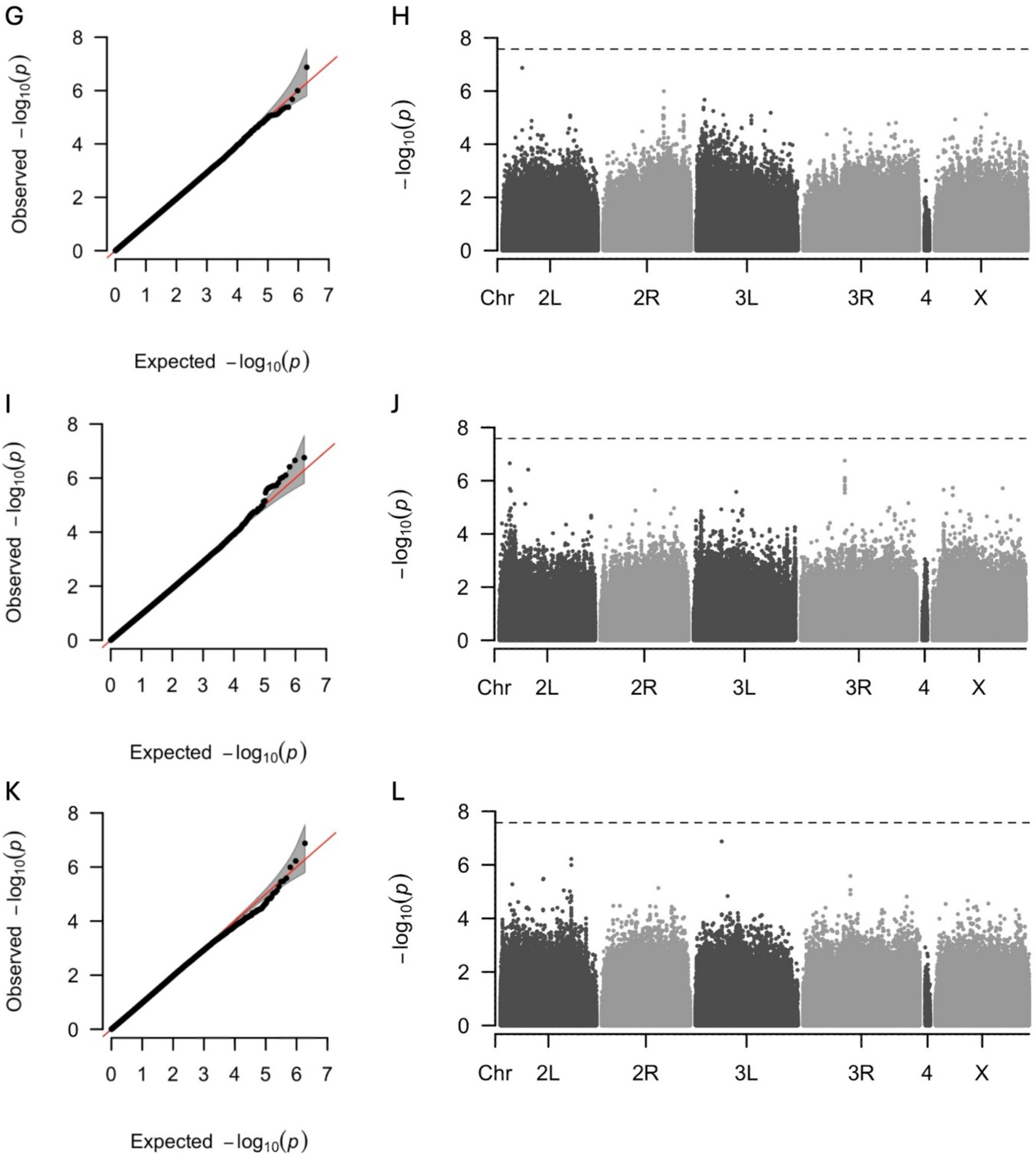


**Figure S6 (continued)**. **Q-Q plot and Manhattan plot for locomotor activity.** Results of a GWAS to identify variants associated with locomotor activity of (**A**, **B**) 7 day-old flies fed the control diet, (**C**, **D**) 7 day-old flies fed the restricted diet, (**E**, **F**) 16 day-old flies fed the control diet, (**G**, **H**) 16 day-old flies fed the restricted diet, (**I**, **J**) genotype-by-diet interaction for 7 day-old flies, (**K**, **L**) genotype-by-diet interaction for 16 day-old flies, (**M**, **N**) genotype-by-age interaction for flies fed the control diet, and (**O**, **P**) genotype-by-age interaction for flies fed the restricted diet. Panels (**A**, **C**, **E**, **G**, **I**, **K, M, O**) are Q-Q plots comparing the observed -log(p) values of each variant to the expected values, with the red line representing the null expectation and the grey area indicating the confidence interval. Panels (**B**, **D**, **F**, **H**, **J**, **L**, **N, P**) are Manhattan plots where each point represents a variant. The y-axis shows the strength of the association between individual variants and dry weight, expressed as -log(p). The dashed horizontal line indicates the significance threshold adjusted for multiple testing using the Bonferroni correction.

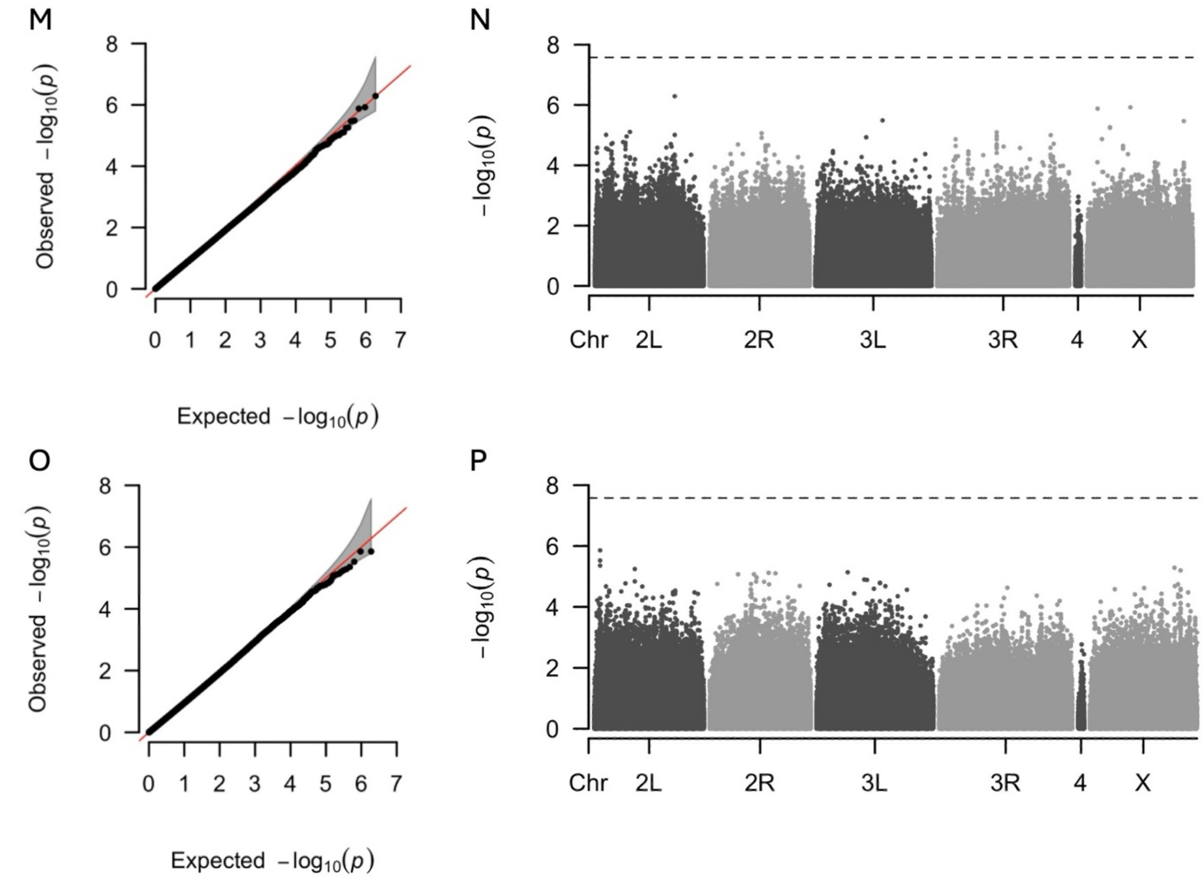


**Figure S6 (continued)**. **Q-Q plot and Manhattan plot for locomotor activity.** Results of a GWAS to identify variants associated with locomotor activity of (**A**, **B**) 7 day-old flies fed the control diet, (**C**, **D**) 7 day-old flies fed the restricted diet, (**E**, **F**) 16 day-old flies fed the control diet, (**G**, **H**) 16 day-old flies fed the restricted diet, (**I**, **J**) genotype-by-diet interaction for 7 day-old flies, (**K**, **L**) genotype-by-diet interaction for 16 day-old flies, (**M**, **N**) genotype-by-age interaction for flies fed the control diet, and (**O**, **P**) genotype-by-age interaction for flies fed the restricted diet. Panels (**A**, **C**, **E**, **G**, **I**, **K, M, O**) are Q-Q plots comparing the observed -log(p) values of each variant to the expected values, with the red line representing the null expectation and the grey area indicating the confidence interval. Panels (**B**, **D**, **F**, **H**, **J**, **L**, **N, P**) are Manhattan plots where each point represents a variant. The y-axis shows the strength of the association between individual variants and dry weight, expressed as -log(p). The dashed horizontal line indicates the significance threshold adjusted for multiple testing using the Bonferroni correction.

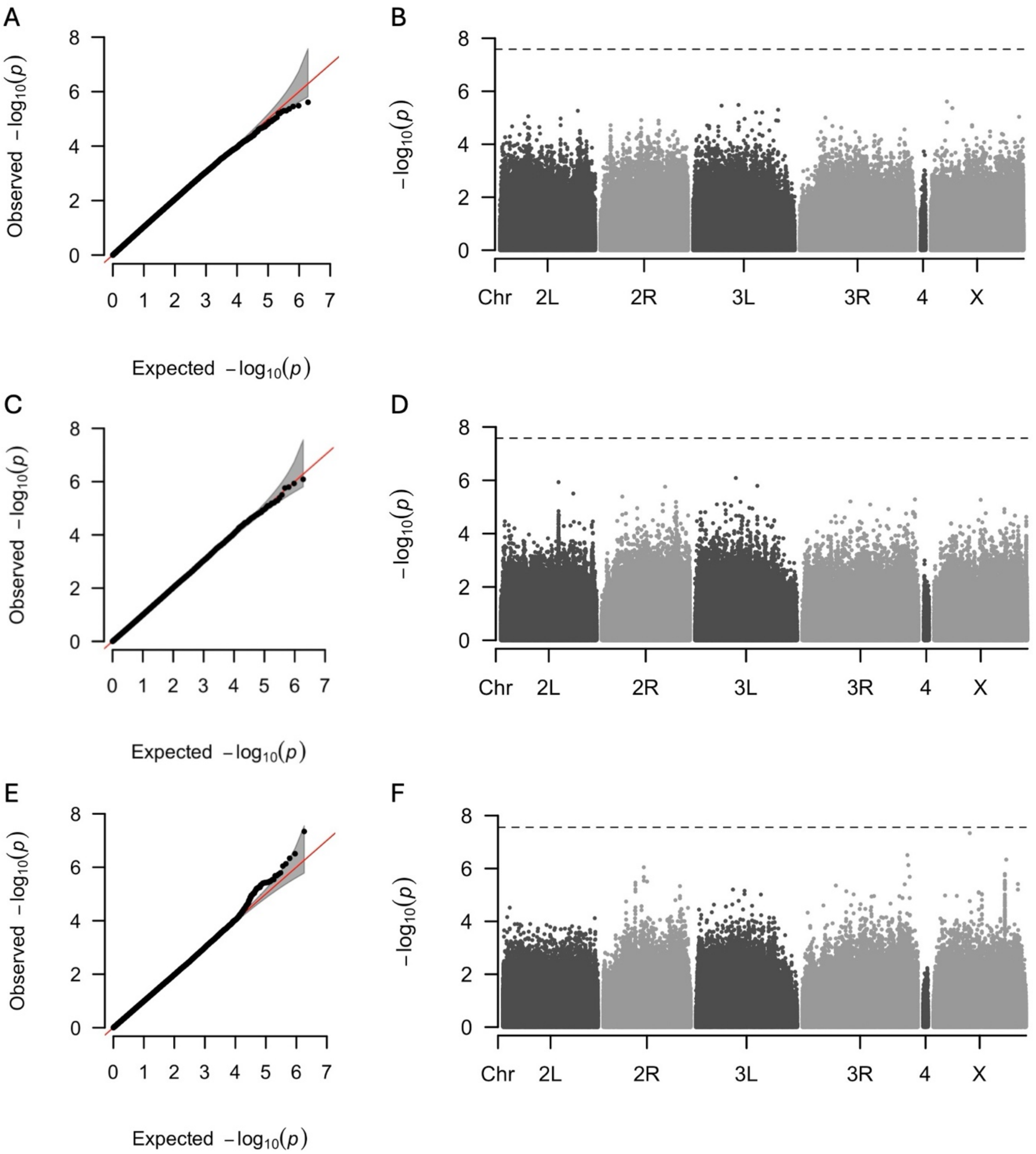


**Figure S7**. **Q-Q plot and Manhattan plot for HKDT.** Results of a GWAS to identify variants associated with HKDT of (**A**, **B**) 7 day-old flies fed the control diet, (**C**, **D**) 7 day-old flies fed the restricted diet, (**E**, **F**) 16 day-old flies fed the restricted diet, (**G**, **H**) genotype-by-diet interaction for 7 day-old flies, (**I**, **J**) genotype-by-diet interaction for 16 day-old flies, (**K**, **L**) genotype-by-age interaction for flies fed the control diet, and (**M**, **N**) genotype-by-age interaction for flies fed the restricted diet. Panels (**A**, **C**, **E**, **G**, **I**, **K, M**are Q-Q plots comparing the observed -log(p) values of each variant to the expected values, with the red line representing the null expectation and the grey area indicating the confidence interval. Panels (**B**, **D**, **F**, **H**, **J**, **L, N**) are Manhattan plots where each point represents a variant. The y-axis shows the strength of the association between individual variants and dry weight, expressed as -log(p). The dashed horizontal line indicates the significance threshold adjusted for multiple testing using the Bonferroni correction.

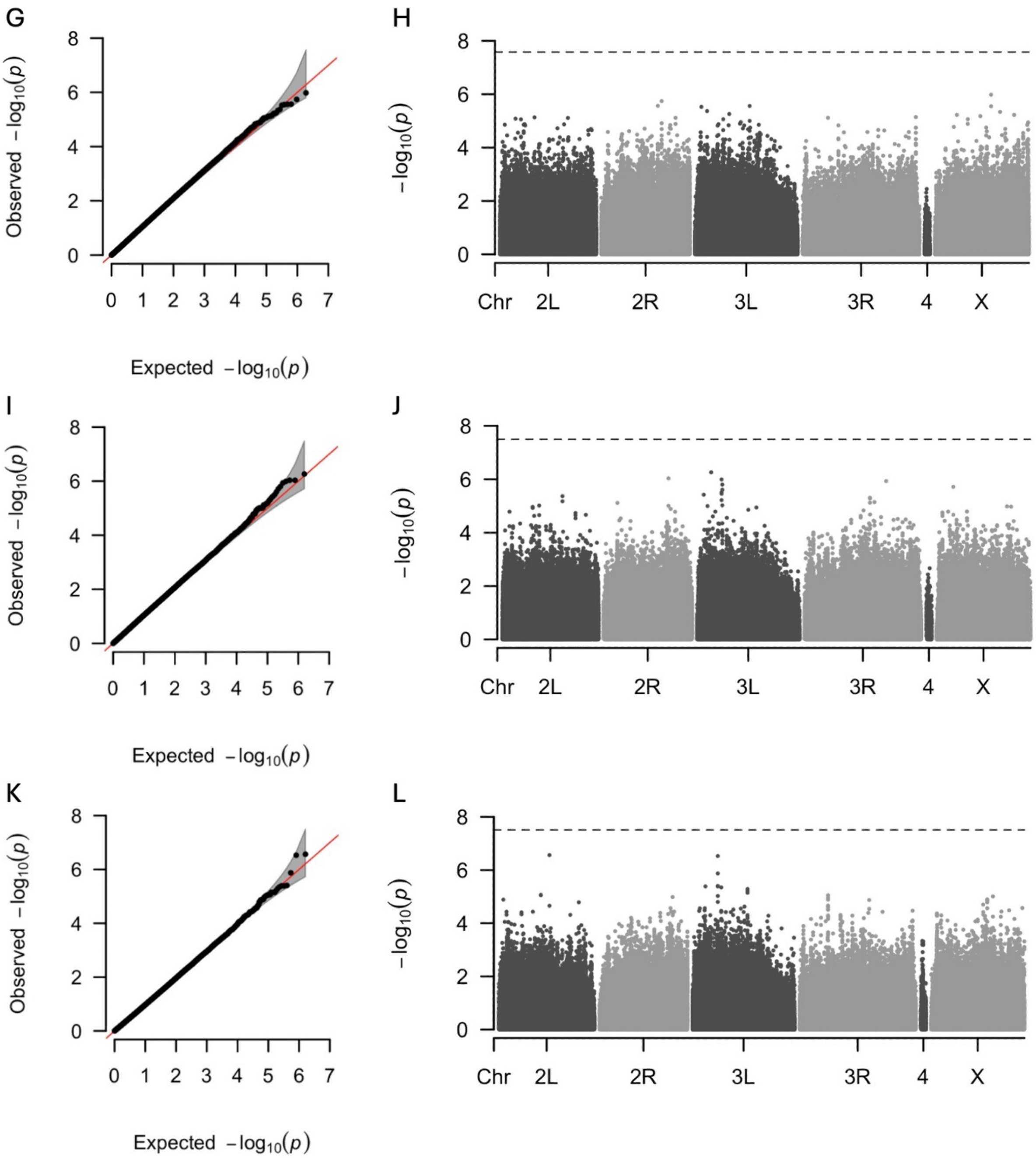


**Figure S7 (continued)**. **Q-Q plot and Manhattan plot for HKDT.** Results of a GWAS to identify variants associated with HKDT of (**A**, **B**) 7 day-old flies fed the control diet, (**C**, **D**) 7 day-old flies fed the restricted diet, (**E**, **F**) 16 day-old flies fed the restricted diet, (**G**, **H**) genotype-by-diet interaction for 7 day-old flies, (**I**, **J**) genotype-by-diet interaction for 16 day-old flies, (**K**, **L**) genotype-by-age interaction for flies fed the control diet, and (**M**, **N**) genotype-by-age interaction for flies fed the restricted diet. Panels (**A**, **C**, **E**, **G**, **I**, **K**, **M**are Q-Q plots comparing the observed -log(p) values of each variant to the expected values, with the red line representing the null expectation and the grey area indicating the confidence interval. Panels (**B**, **D**, **F**, **H**, **J**, **L**, **N**) are Manhattan plots where each point represents a variant. The y-axis shows the strength of the association between individual variants and dry weight, expressed as -log(p). The dashed horizontal line indicates the significance threshold adjusted for multiple testing using the Bonferroni correction.

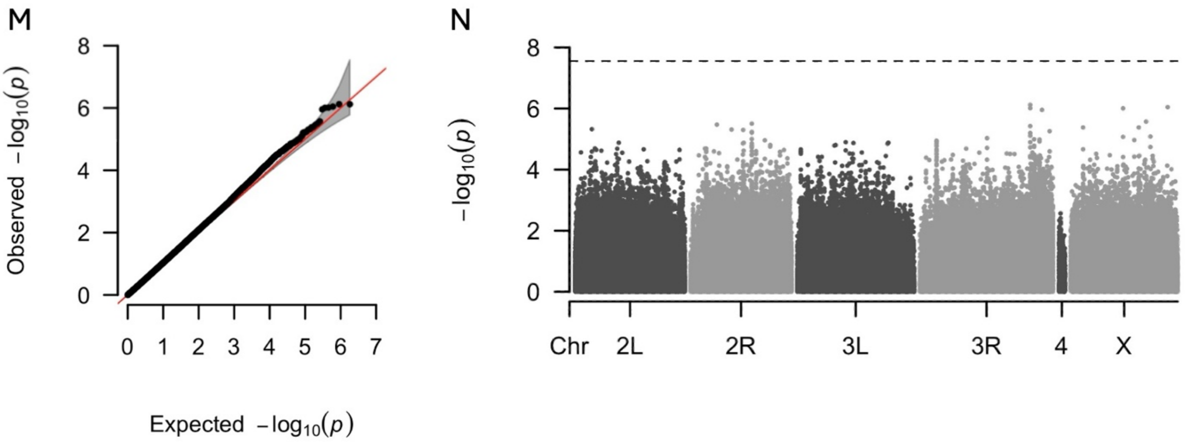


**Figure S7 (continued)**. **Q-Q plot and Manhattan plot for HKDT.** Results of a GWAS to identify variants associated with HKDT of (**A**, **B**) 7 day-old flies fed the control diet, (**C**, **D**) 7 day-old flies fed the restricted diet, (**E**, **F**) 16 day-old flies fed the restricted diet, (**G**, **H**) genotype-by-diet interaction for 7 day-old flies, (**I**, **J**) genotype-by-diet interaction for 16 day-old flies, (**K**, **L**) genotype-by-age interaction for flies fed the control diet, and (**M**, **N**) genotype-by-age interaction for flies fed the restricted diet. Panels (**A**, **C**, **E**, **G**, **I**, **K, M**are Q-Q plots comparing the observed -log(p) values of each variant to the expected values, with the red line representing the null expectation and the grey area indicating the confidence interval. Panels (**B**, **D**, **F**, **H**, **J**, **L, N**) are Manhattan plots where each point represents a variant. The y-axis shows the strength of the association between individual variants and dry weight, expressed as -log(p). The dashed horizontal line indicates the significance threshold adjusted for multiple testing using the Bonferroni correction.

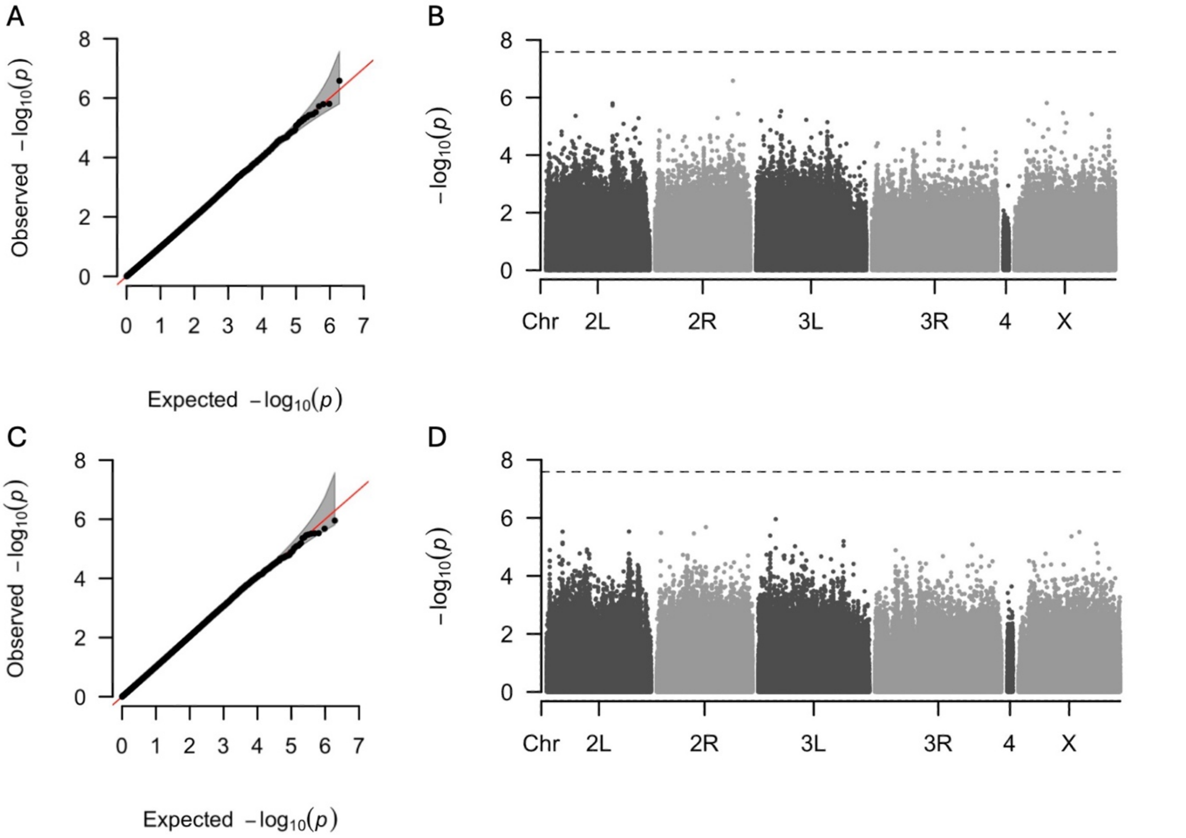


**Figure S8**. **Q-Q plot and Manhattan plot for lifespan.** Results of a GWAS to identify variants associated with lifespan of (**A**, **B**) flies fed the control diet and (**C**, **D**) flies fed the restricted diet. Panels (**A**, **C**) are Q-Q plots comparing the observed -

log(p) values of each variant to the expected values, with the red line representing the null expectation and the grey area indicating the confidence interval. Panels (**B**, **D**) are Manhattan plots where each point represents a variant. The y-axis shows the strength of the association between individual variants and dry weight, expressed as -log(p). The dashed horizontal line indicates the significance threshold adjusted for multiple testing using the Bonferroni correction.

**Table S1.** Number of flies across age groups and diets for both unbalanced and balanced datasets of dry weight measurements.

| Type of test | Unbalanced | | Balanced | |
|---|---|---|---|---|
| | Control diet | Restricted diet | Control diet | Restricted diet |
| Two-way age 1 | 778 | 783 | 778 | 783 |
| Two-way age 2 | 756 | 748 | 740 | 740 |
| Two-way age 3 | 785 | 655 | 718 | 642 |
| Two-way age 4 | 998 | 128 | 308 | 128 |
| Two-way age 5 | 521 | 9 | 32 | 9 |
| Three-way age 1 and 2 | 1534 | 1531 | 1479 | 1483 |
| Three-way age 1, 2 and 3 | 2319 | 2186 | 2030 | 1963 |
| Three-way age 1, 2, 3 and 4 | 3317 | 2314 | 834 | 650 |

**Table S2.** Number of flies across age groups and diets for both unbalanced and balanced datasets of locomotor activity measurements.

| Type of test | Unbalanced | | Balanced | |
|---|---|---|---|---|
| | Control diet | Restricted diet | Control diet | Restricted diet |
| Two-way age 1 | 770 | 758 | 770 | 758 |
| Two-way age 2 | 617 | 672 | 602 | 660 |
| Two-way age 3 | 618 | 557 | 579 | 541 |
| Two-way age 4 | 900 | 87 | 235 | 86 |
| Two-way age 5 | 489 | 8 | 25 | 8 |
| Three-way age 1 and 2 | 1387 | 1430 | 1320 | 1367 |
| Three-way age 1, 2 and 3 | 2005 | 1987 | 1755 | 1749 |
| Three-way age 1, 2, 3 and 4 | 2905 | 2074 | 623 | 481 |

**Table S3.** Number of flies across age groups and diets for both unbalanced and balanced datasets of HKDT measurements.

| Type of test | Unbalanced | | Balanced | |
|---|---|---|---|---|
| | Control diet | Restricted diet | Control diet | Restricted diet |
| Two-way age 1 | 691 | 558 | 669 | 558 |
| Two-way age 2 | 225 | 430 | 214 | 322 |
| Two-way age 3 | 228 | 320 | 257 | 261 |
| Two-way age 4 | 553 | 43 | 105 | 39 |
| Two-way age 5 | 430 | 8 | 17 | 8 |
| Three-way age 1 and 2 | 916 | 988 | 645 | 696 |
| Three-way age 1, 2 and 3 | 1204 | 1308 | 714 | 748 |
| Three-way age 1, 2, 3 and 4 | 1757 | 1351 | 194 | 140 |

**Table S4.** Summary of full and reduced ANOVA models assessing dry weight of DGRP flies at 7, 16 and 25 days old and fed control and restricted diets. Diet and age were considered fixed effects, and DGRP line is a random effect.

| Analysis | Source | DF | MS | F Value | *p*-value | $\sigma^2$ | Lower | Upper | $H^2$ |
|---|---|---|---|---|---|---|---|---|---|
| 7 days old on control diet | L | 97 | 6.578E-3 | 5.03 | <0.0001 | 0.686E-3 | 0.442E-3 | 0.929E-3 | 0.3437 |
| | Rep(L) | 170 | 1.308E-3 | 1.00 | 0.4956 | -0.594E-6 | -0.11E-3 | 0.106E-3 | |
| | $\epsilon$ | 510 | 1.310E-3 | - | - | 1.310E-3 | 1.163E-3 | 0.149E-3 | |
| 7 days old on restricted diet | L | 97 | 3.885E-3 | 2.96 | <0.0001 | 0.332E-3 | 0.185E-3 | 0.478E-3 | 0.2542 |
| | Rep(L) | 169 | 1.267E-3 | 1.30 | 0.0155 | 0.107E-3 | -3.03E-6 | 0.217E-3 | |
| | $\epsilon$ | 516 | 0.974E-3 | - | - | 0.974E-3 | 0.866E-3 | 1.105E-3 | |
| 7 days old on both diets | D | 1 | 0.0761 | 26.61 | <0.0001 | Fixed | Fixed | Fixed | 0.3081 |
| | L | 97 | 7.572E-3 | 2.63 | <0.0001 | 0.304E-3 | 0.156E-3 | 0.452E-3 | |
| | D×L | 97 | 2.880E-3 | 2.20 | <0.0001 | 0.204E-3 | 0.092E-3 | 0.315E-3 | |
| | Rep(D×L) | 339 | 1.288E-3 | 1.13 | 0.0823 | 0.054E-3 | -0.02E-3 | 0.131E-3 | |
| | $\epsilon$ | 1026 | 1.141E-3 | - | - | 1.141E-3 | 1.049E-3 | 1.247E-3 | |
| 16 days old on control diet | L | 94 | 3.643E-3 | 3.47 | <0.0001 | 0.336E-3 | 0.196E-3 | 0.475E-3 | 0.2451 |
| | Rep(L) | 167 | 1.049E-3 | 1.01 | 0.4472 | 4.417E-6 | -0.008E-3 | 0.093E-3 | |
| | $\epsilon$ | 494 | 1.035E-3 | - | - | 1.035E-3 | 0.917E-3 | 1.177E-3 | |

| | | | | | | | | | |
|---|---|---|---|---|---|---|---|---|---|
| 16 days old on restricted diet | L | 93 | 4.386E-3 | 3.68 | <0.0001 | 0.420E-3 | 0.252E-3 | 0.589E-3 | 0.2793 |
| | Rep(L) | 164 | 1.179E-3 | 1.09 | 0.2473 | 0.035E-3 | -0.007E-3 | 0.137E-3 | |
| | $\epsilon$ | 490 | 1.084E-3 | - | - | 1.084E-3 | 0.960E-3 | 1.233E-3 | |
| 16 days old on both diets | D | 1 | 0.0560 | 31.79 | <0.0001 | Fixed | Fixed | Fixed | 0.2615 |
| | L | 95 | 6.134E-3 | 3.46 | <0.0001 | 0.290E-3 | 0.169E-3 | 0.412E-3 | |
| | D×L | 92 | 1.770E-3 | 1.58 | 0.0026 | 0.085E-3 | 0.015E-3 | 0.115E-3 | |
| | Rep(D×L) | 331 | 1.113E-3 | 1.05 | 0.2832 | 0.020E-3 | -0.005E-3 | 0.085E-3 | |
| | $\epsilon$ | 984 | 1.059E-3 | - | - | 1.059E-3 | 0.971E-3 | 1.159E-3 | |
| 7 and 16 days old on control diet | A | 1 | 0.949E-3 | 0.45 | 0.5036 | Fixed | Fixed | Fixed | 0.3043 |
| | L | 97 | 8.058E-3 | 3.82 | <0.0001 | 0.393E-3 | 0.236E-3 | 0.549E-3 | |
| | A×L | 94 | 2.110E-3 | 1.79 | 0.0002 | 0.121E-3 | 0.038E-3 | 0.203E-3 | |
| | Rep(A×L) | 337 | 1.180E-3 | 1.00 | 0.4731 | 2.039E-6 | -0.006E-3 | 0.069E-3 | |
| | $\epsilon$ | 1004 | 1.175E-3 | - | - | 1.175E-3 | 1.078E-3 | 1.284E-3 | |
| 7 and 16 days old on restricted diet | A | 1 | 2.430E-3 | 1.12 | 0.2929 | Fixed | Fixed | Fixed | 0.2657 |
| | L | 97 | 5.933E-3 | 2.69 | <0.0001 | 0.248E-3 | 0.128E-3 | 0.369E-3 | |
| | A×L | 93 | 2.194E-3 | 1.76 | 0.0002 | 0.124E-3 | 0.042E-3 | 0.205E-3 | |
| | Rep(A×L) | 333 | 1.223E-3 | 1.19 | 0.0232 | 0.072E-3 | -3.7E-6 | 0.148E-3 | |
| | $\epsilon$ | 1006 | 1.028E-3 | - | - | 1.028E-3 | 0.943E-3 | 1.124E-3 | |
| 7 and 16 days old on both diets | A | 1 | 2.748E-3 | 1.35 | 0.2489 | Fixed | Fixed | Fixed | 0.2928 |
| | D | 1 | 0.1362 | 57.72 | <0.0001 | Fixed | Fixed | Fixed | |
| | L | 97 | 0.0116 | 5.41 | <0.0001 | 0.314E-3 | 0.199E-3 | 0.429E-3 | |
| | A×D | 1 | 0.191E-3 | 0.08 | 0.7723 | Fixed | Fixed | Fixed | |
| | A×L | 95 | 2.048E-3 | 0.90 | 0.6941 | -0.002E-3 | -0.07E-3 | 0.041E-3 | |
| | D×L | 97 | 2.368E-3 | 1.04 | 0.4271 | 5.948E-6 | -0.06E-3 | 0.068E-3 | |
| | A×D×L | 92 | 2.275E-3 | 1.88 | <0.0001 | 0.138E-3 | 0.050E-3 | 0.227E-3 | |
| | Rep(A×D×L) | 670 | 1.202E-3 | 1.09 | 0.0800 | 0.037E-3 | -0.01E-3 | 0.087E-3 | |
| | $\epsilon$ | 2010 | 1.101E-3 | - | - | 1.101E-3 | 1.036E-3 | 1.172E-3 | |
| 25 days old on control diet | L | 94 | 3.751E-3 | 2.87 | <0.0001 | 0.306E-3 | 0.165E-3 | 0.448E-3 | 0.2546 |
| | Rep(L) | 153 | 1.289E-3 | 1.44 | 0.0018 | 0.130E-3 | 0.232E-3 | 0.232E-3 | |
| | $\epsilon$ | 537 | 0.896E-3 | - | - | 0.896E-3 | 1.014E-3 | 1.014E-3 | |
| 25 days old on restricted diet | L | 86 | 3.105E-3 | 2.10 | <0.0001 | 0.224E-3 | 0.088E-3 | 0.361E-3 | 0.1435 |
| | Rep(L) | 125 | 1.470E-3 | 1.10 | 0.2433 | 0.045E-3 | -0.09E-3 | 0.182E-3 | |
| | $\epsilon$ | 443 | 1.337E-3 | - | - | 1.337E-3 | 1.177E-3 | 1.532E-3 | |
| 25 days old on both diets | D | 1 | 0.3638 | 23.28 | <0.0001 | Fixed | Fixed | Fixed | 0.1949 |
| | L | 96 | 4.744E-3 | 2.87 | <0.0001 | 0.226E-3 | 0.120E-3 | 0.331E-3 | |
| | D×L | 84 | 1.654E-3 | 1.21 | 0.1329 | 0.039E-3 | -0.04E-3 | 0.116E-3 | |
| | Rep(A×D×L) | 278 | 1.370E-3 | 1.25 | 0.0083 | 0.092E-3 | 8.871E-6 | 0.175E-3 | |
| | $\epsilon$ | 980 | 1.095E-3 | - | - | 1.095E-3 | 1.005E-3 | 1.199E-3 | |
| 7, 16 and 25 days old on both diets | A | 2 | 6.195E-3 | 3.13 | 0.0460 | Fixed | Fixed | Fixed | 0.2594 |
| | D | 1 | 0.1611 | 72.88 | <0.0001 | Fixed | Fixed | Fixed | |
| | L | 97 | 0.0136 | 5.87 | <0.0001 | 0.267E-3 | 0.175E-3 | 0.359E-3 | |
| | A×D | 2 | 0.393E-3 | 0.20 | 0.8204 | Fixed | Fixed | Fixed | |
| | A×L | 191 | 2.050E-3 | 1.01 | 0.4820 | 0.989E-6 | -0.04E-3 | 0.041E-3 | |
| | D×L | 97 | 2.289E-3 | 1.13 | 0.2408 | 0.013E-3 | -0.02E-3 | 0.050E-3 | |
| | A×D×L | 176 | 2.039E-3 | 1.62 | <0.0001 | 0.104E-3 | 0.044E-3 | 0.165E-3 | |
| | Rep(A×D×L) | 948 | 1.251E-3 | 1.14 | 0.0064 | 0.054E-3 | 0.011E-3 | 0.097E-3 | |
| | $\epsilon$ | 2990 | 1.099E-3 | - | - | 1.099E-3 | 1.046E-3 | 1.157E-3 | |

**Table S5.** Summary of full and reduced ANOVA models assessing locomtor activity of DGRP flies at 7, 16 and 25 days old and fed control and restricted diets. Diet and age were considered fixed effects, and DGRP line is a random effect.

| Analysis | Source | DF | MS | F Value | *p*-value | $\sigma^2$ | Lower | Upper | $H^2$ |
|---|---|---|---|---|---|---|---|---|---|
| 7 days old on control diet | L | 97 | 3.7961 | 5.21 | <0.0001 | 0.4034 | 0.2627 | 0.5442 | 0.3717 |
| | Rep(L) | 170 | 0.7225 | 1.06 | 0.3143 | 0.0152 | -0.0505 | 0.0809 | |
| | $\epsilon$ | 502 | 0.6819 | - | - | 0.6819 | 0.6048 | 0.7748 | |
| 7 days old on restricted diet | L | 97 | 5.3835 | 5.11 | <0.0001 | 0.5805 | 0.3756 | 0.7855 | 0.4221 |
| | Rep(L) | 168 | 1.0182 | 1.28 | 0.0219 | 0.0848 | -0.0055 | 0.1750 | |
| | $\epsilon$ | 492 | 0.7949 | - | - | 0.7949 | 0.7042 | 0.9044 | |
| 7 days old on both diets | D | 1 | 0.0010 | 0.00 | 0.9807 | Fixed | Fixed | Fixed | 0.4004 |
| | L | 97 | 7.4081 | 4.20 | <0.0001 | 0.3762 | 0.2328 | 0.5197 | |
| | D×L | 97 | 1.7623 | 1.98 | <0.0001 | 0.1164 | 0.0488 | 0.1841 | |
| | Rep(D×L) | 338 | 0.8695 | 1.18 | 0.0299 | 0.0496 | -0.0051 | 0.1043 | |
| | $\epsilon$ | 994 | 0.7378 | - | - | 0.7378 | 0.6770 | 0.8073 | |
| | L | 92 | 6.3577 | 5.14 | <0.0001 | 0.8203 | 0.5191 | 1.1215 | 0.4533 |

| | | | | | | | | | |
|---|---|---|---|---|---|---|---|---|---|
| 16 days old on control diet | Rep(L) | 144 | 1.2140 | 1.23 | 0.0642 | 0.0933 | -0.0399 | 0.2265 | |
| | $\epsilon$ | 380 | 0.9892 | - | - | 0.9892 | 0.8624 | 1.1465 | |
| 16 days old on restricted diet | L | 93 | 4.9050 | 5.84 | <0.0001 | 0.6087 | 0.4005 | 0.8169 | 0.4006 |
| | Rep(L) | 161 | 0.8474 | 0.93 | 0.7010 | -0.0263 | -0.1091 | 0.0565 | |
| | $\epsilon$ | 417 | 0.9108 | - | - | 0.9108 | 0.7988 | 1.0483 | |
| 16 days old on both diets | D | 1 | 34.0410 | 13.64 | 0.0004 | Fixed | Fixed | Fixed | 0.4315 |
| | L | 95 | 8.2341 | 3.09 | <0.0001 | 0.4574 | 0.2560 | 0.6588 | |
| | D×L | 90 | 2.6653 | 2.60 | <0.0001 | 0.2622 | 0.1345 | 0.3899 | |
| | Rep(D×L) | 305 | 1.0205 | 1.08 | 0.2150 | 0.0300 | -0.0464 | 0.1063 | |
| | $\epsilon$ | 797 | 0.9482 | - | - | 0.9482 | 0.8616 | 1.0487 | |
| 7 and 16 days old on control diet | A | 1 | 16.944 | 4.57 | 0.0351 | Fixed | Fixed | Fixed | 0.4281 |
| | L | 97 | 6.1006 | 1.53 | 0.0200 | 0.1616 | 0.0088 | 0.3144 | |
| | A×L | 92 | 3.9510 | 4.12 | <0.0001 | 0.4479 | 0.2735 | 0.6222 | |
| | Rep(A×L) | 314 | 0.9479 | 1.16 | 0.0477 | 0.0524 | -0.0156 | 0.1204 | |
| | $\epsilon$ | 882 | 0.8143 | - | - | 0.8143 | 0.7433 | 0.8960 | |
| 7 and 16 days old on restricted diet | A | 1 | 1.1681 | 0.60 | 0.4417 | Fixed | Fixed | Fixed | 0.4126 |
| | L | 97 | 8.0454 | 3.98 | <0.0001 | 0.4417 | 0.2691 | 0.6143 | |
| | A×L | 93 | 2.0150 | 2.14 | <0.0001 | 0.1541 | 0.0714 | 0.2368 | |
| | Rep(A×L) | 329 | 0.9346 | 1.10 | 0.1380 | 0.0343 | -0.0263 | 0.0948 | |
| | $\epsilon$ | 909 | 0.8481 | - | - | 0.8481 | 0.7752 | 0.9318 | |
| 7 and 16 days old on both diets | A | 1 | 4.5364 | 1.38 | 0.2433 | Fixed | Fixed | Fixed | 0.4227 |
| | D | 1 | 18.8916 | 10.46 | 0.0016 | Fixed | Fixed | Fixed | |
| | L | 97 | 12.0457 | 4.42 | <0.0001 | 0.3559 | 0.2177 | 0.4941 | |
| | A×D | 1 | 17.4111 | 7.01 | 0.0095 | Fixed | Fixed | Fixed | |
| | A×L | 95 | 3.4283 | 1.33 | 0.0848 | 0.0656 | -0.0261 | 0.1573 | |
| | D×L | 97 | 1.8744 | 0.72 | 0.9413 | -0.0553 | -0.1224 | 0.0118 | |
| | A×D×L | 90 | 2.5778 | 2.71 | <0.0001 | 0.2426 | 0.1282 | 0.3571 | |
| | Rep(A×D×L) | 643 | 0.9411 | 1.13 | 0.0266 | 0.0432 | -0.0026 | 0.0890 | |
| | $\epsilon$ | 1791 | 0.8315 | - | - | 0.8315 | 0.7796 | 0.8887 | |
| 25 days old on control diet | L | 91 | 5.8282 | 3.58 | <0.0001 | 0.6632 | 0.3847 | 0.9418 | 0.4159 |
| | Rep(L) | 134 | 1.6315 | 1.75 | <0.0001 | 0.2665 | 0.1154 | 0.4176 | |
| | $\epsilon$ | 392 | 0.9313 | - | - | 0.9313 | 0.8135 | 1.0767 | |
| 25 days old on restricted diet | L | 86 | 2.8888 | 2.77 | <0.0001 | 0.3037 | 0.1517 | 0.4556 | 0.2380 |
| | Rep(L) | 119 | 1.0438 | 1.07 | 0.3084 | 0.0275 | -0.0808 | 0.1359 | |
| | $\epsilon$ | 351 | 0.9722 | - | - | 0.9722 | 0.8430 | 1.1337 | |
| 25 days old on both diets | D | 1 | 0.9845 | 0.55 | 0.4583 | Fixed | Fixed | Fixed | 0.3446 |
| | L | 95 | 6.1768 | 3.19 | <0.0001 | 0.3929 | 0.2112 | 0.5746 | |
| | D×L | 82 | 1.9615 | 1.47 | 0.0125 | 0.1070 | -0.0091 | 0.2230 | |
| | Rep(A×D×L) | 253 | 1.3551 | 1.43 | 0.0002 | 0.1547 | 0.0629 | 0.2465 | |
| | $\epsilon$ | 743 | 0.9506 | - | - | 0.9506 | 0.8609 | 1.0552 | |
| 7, 16 and 25 days old on both diets | A | 2 | 18.6262 | 6.22 | 0.0024 | Fixed | Fixed | Fixed | 0.3931 |
| | D | 1 | 14.7378 | 7.73 | 0.0064 | Fixed | Fixed | Fixed | |
| | L | 97 | 13.5681 | 4.33 | <0.0001 | 0.3003 | 0.1906 | 0.4101 | |
| | A×D | 2 | 9.9269 | 4.91 | 0.0084 | Fixed | Fixed | Fixed | |
| | A×L | 190 | 3.2651 | 1.54 | 0.0020 | 0.0945 | 0.0279 | 0.1611 | |
| | D×L | 97 | 2.0550 | 0.97 | 0.5586 | -0.0036 | -0.0416 | 0.0345 | |
| | A×D×L | 172 | 2.1466 | 2.02 | <0.0001 | 0.1700 | 0.0956 | 0.2444 | |
| | Rep(A×D×L) | 896 | 1.0580 | 1.22 | 0.0001 | 0.0749 | 0.0337 | 0.1160 | |
| | $\epsilon$ | 2534 | 0.8664 | - | - | 0.8664 | 0.8206 | 0.9162 | |

**Table S6.** Summary of full and reduced ANOVA models assessing HKDT of DGRP flies at 7, 16 and 25 days old and fed control and restricted diets. Diet and age were considered fixed effects, and DGRP line is a random effect.

| Analysis | Source | DF | MS | F Value | *p*-value | $\sigma^2$ | Lower | Upper | $H^2$ |
|---|---|---|---|---|---|---|---|---|---|
| 7 days old on control diet | L | 96 | 531.8690 | 4.23 | <0.0001 | 59.3788 | 36.9663 | 81.7913 | 0.3865 |
| | Rep(L) | 157 | 121.8875 | 1.29 | 0.0222 | 10.9662 | -2.5497 | 24.4821 | |
| | $\epsilon$ | 437 | 94.2677 | - | - | 94.2677 | 82.9184 | 108.13 | |
| 7 days old on restricted diet | L | 93 | 394.1051 | 3.60 | <0.0001 | 51.3928 | 28.8399 | 73.9457 | 0.3264 |
| | Rep(L) | 138 | 109.0432 | 1.03 | 0.4159 | 1.3612 | -13.0343 | 15.7566 | |
| | $\epsilon$ | 325 | 106.0603 | - | - | 106.06 | 91.4711 | 124.46 | |
| 7 days old on both diets | D | 1 | 16006 | 58.94 | <0.0001 | Fixed | Fixed | Fixed | 0.3613 |
| | L | 96 | 608.4325 | 2.06 | 0.0003 | 26.5633 | 9.7362 | 43.3904 | |
| | D×L | 93 | 292.9536 | 2.50 | <0.0001 | 29.6048 | 15.4089 | 43.8007 | |

| | | | | | | | | | |
|---|---|---|---|---|---|---|---|---|---|
| | Rep(D×L) | 295 | 115.8790 | 1.17 | 0.0522 | 7.0095 | -2.8255 | 16.8446 | |
| | $\epsilon$ | 762 | 99.2973 | - | - | 99.2973 | 90.0350 | 110.07 | |
| 16 days old on control diet | L | 61 | 229.8621 | 1.35 | 0.1267 | 17.6157 | -15.5556 | 50.7870 | 0.1137 |
| | Rep(L) | 57 | 170.3693 | 1.24 | 0.1695 | 18.2846 | -36.9643 | 73.5335 | |
| | $\epsilon$ | 106 | 137.3836 | - | - | 137.38 | 106.78 | 183.41 | |
| 16 days old on restricted diet | L | 83 | 250.4292 | 3.05 | <0.0001 | 35.6428 | 16.2972 | 54.9883 | 0.3130 |
| | Rep(L) | 114 | 81.9705 | 1.05 | 0.3795 | 1.8777 | -15.4180 | 19.1733 | |
| | $\epsilon$ | 232 | 78.2379 | - | - | 78.2379 | 65.7461 | 94.6823 | |
| 16 days old on both diets | D | 1 | 1470.7298 | 7.26 | 0.0088 | Fixed | Fixed | Fixed | 0.2475 |
| | L | 86 | 230.6495 | 0.98 | 0.5364 | -0.6597 | -18.6607 | 17.3413 | |
| | D×L | 58 | 227.3544 | 2.05 | 0.0002 | 32.4862 | 4.2173 | 60.7551 | |
| | Rep(D×L) | 171 | 111.4368 | 1.15 | 0.1393 | 7.6041 | -12.1045 | 27.3127 | |
| | $\epsilon$ | 338 | 96.7865 | - | - | 96.7865 | 83.7044 | 113.21 | |
| 7 and 16 days old on control diet | A | 1 | 34808 | 181.80 | <0.0001 | Fixed | Fixed | Fixed | 0.3512 |
| | L | 96 | 518.1841 | 2.21 | 0.0013 | 36.0583 | 10.3376 | 61.7790 | |
| | A×L | 61 | 212.7383 | 1.62 | 0.0054 | 19.5184 | -0.6681 | 39.7049 | |
| | Rep(A×L) | 214 | 134.8009 | 1.31 | 0.0073 | 13.7940 | -0.8023 | 28.3903 | |
| | $\epsilon$ | 543 | 102.6844 | - | - | 102.68 | 91.4866 | 116.08 | |
| 7 and 16 days old on restricted diet | A | 1 | 6256.9640 | 35.56 | <0.0001 | Fixed | Fixed | Fixed | 0.3293 |
| | L | 94 | 440.3188 | 2.33 | <0.0001 | 27.4449 | 10.2982 | 44.5916 | |
| | A×L | 82 | 189.9159 | 1.96 | <0.0001 | 18.9446 | 6.3002 | 31.5890 | |
| | Rep(A×L) | 252 | 96.7960 | 1.02 | 0.4051 | 1.1071 | -9.4287 | 11.6430 | |
| | $\epsilon$ | 557 | 94.4718 | - | - | 94.4718 | 84.2890 | 106.63 | |
| 7 and 16 days old on both diets | A | 1 | 34579 | 207.86 | <0.0001 | Fixed | Fixed | Fixed | 0.3234 |
| | D | 1 | 1130.5781 | 4.83 | 0.0298 | Fixed | Fixed | Fixed | |
| | L | 96 | 558.1189 | 2.18 | 0.0157 | 19.4587 | 6.4627 | 32.4547 | |
| | A×D | 1 | 7162.0576 | 36.57 | <0.0001 | Fixed | Fixed | Fixed | |
| | A×L | 86 | 182.3947 | 0.87 | 0.7284 | -3.8503 | -14.3266 | 6.6260 | |
| | D×L | 94 | 286.7512 | 1.31 | 0.1426 | 9.1820 | -4.7773 | 23.1412 | |
| | A×D×L | 57 | 210.4926 | 1.85 | 0.0003 | 22.3075 | 5.4590 | 39.1559 | |
| | Rep(A×D×L) | 466 | 114.2489 | 1.16 | 0.0273 | 7.1323 | -1.6360 | 15.9005 | |
| | $\epsilon$ | 1100 | 98.5258 | - | - | 98.5258 | 90.7837 | 107.31 | |
| 25 days old on control diet | L | 69 | 279.3379 | 2.47 | <0.0001 | 44.1435 | 16.9072 | 71.3799 | 0.2263 |
| | Rep(L) | 67 | 110.9311 | 151 | 0.9224 | -19.8269 | -40.4556 | 0.8018 | |
| | $\epsilon$ | 151 | 150.9137 | - | - | 150.91 | 121.92 | 191.70 | |
| 25 days old on restricted diet | L | 73 | 254.5007 | 1.44 | 0.0577 | 19.0153 | -1.6391 | 39.6697 | 0.2059 |
| | Rep(L) | 79 | 177.7359 | 2.42 | <0.0001 | 52.3019 | 20.3548 | 84.2490 | |
| | $\epsilon$ | 167 | 73.3219 | - | - | 73.3219 | 59.8254 | 91.9893 | |
| 25 days old on both diets | D | 1 | 414.8376 | 1.77 | 0.1874 | Fixed | Fixed | Fixed | 0.2257 |
| | L | 84 | 259.2257 | 1.00 | 0.5112 | -0.1370 | -20.7342 | 20.4603 | |
| | D×L | 58 | 262.7501 | 1.81 | 0.0020 | 32.2515 | 6.8154 | 57.6875 | |
| | Rep(A×D×L) | 146 | 147.0789 | 1.34 | 0.0182 | 18.4046 | -2.0600 | 38.8692 | |
| | $\epsilon$ | 318 | 110.1658 | - | - | 110.17 | 94.8639 | 129.51 | |
| 7, 16 and 25 days old on both diets | A | 2 | 25938 | 140.32 | <0.0001 | Fixed | Fixed | Fixed | 0.2974 |
| | D | 1 | 189.4276 | 0.75 | 0.3870 | Fixed | Fixed | Fixed | |
| | L | 96 | 546.6994 | 1.74 | 0.0143 | 11.9537 | 1.5929 | 22.3146 | |
| | A×D | 2 | 5102.5452 | 26.87 | <0.0001 | Fixed | Fixed | Fixed | |
| | A×L | 170 | 206.2055 | 1.02 | 0.4686 | 0.4400 | -8.4650 | 9.3450 | |
| | D×L | 94 | 309.9370 | 1.51 | 0.0165 | 11.3762 | 1.5052 | 21.2471 | |
| | A×D×L | 115 | 205.9576 | 1.70 | <0.0001 | 20.0453 | 7.3953 | 32.6954 | |
| | Rep(A×D×L) | 612 | 122.0809 | 1.21 | 0.0026 | 9.7098 | 1.4948 | 17.9248 | |
| | $\epsilon$ | 1418 | 101.1362 | - | - | 101.14 | 94.0859 | 109.01 | |

**Table S7.** Summary of full and reduced ANOVA models for balanced data assessing dry weight of DGRP flies at 7, 16 and 25 days old and fed control and restricted diets. Diet and age were considered fixed effects, and DGRP line is a random effect.

| Analysis | Source | DF | MS | F Value | *p*-value | $\sigma^2$ | Lower | Upper | $H^2$ |
|---|---|---|---|---|---|---|---|---|---|
| 7 days old on control diet | L | 97 | 6.578E-3 | 5.03 | <0.0001 | 0.686E-3 | 0.442E-3 | 0.929E-3 | 0.3437 |
| | Rep(L) | 170 | 1.308E-3 | 1.00 | 0.4956 | -0.594E-6 | -0.11E-3 | 0.106E-3 | |
| | $\epsilon$ | 510 | 1.310E-3 | - | - | 1.310E-3 | 1.163E-3 | 1.487E-3 | |
| 7 days old on restricted diet | L | 97 | 3.885E-3 | 2.96 | <0.0001 | 0.332E-3 | 0.185E-3 | 0.478E-3 | 0.2542 |
| | Rep(L) | 169 | 1.267E-3 | 1.30 | 0.0155 | 0.107E-3 | -3.03E-6 | 0.217E-3 | |
| | $\epsilon$ | 516 | 0.974E-3 | - | - | 0.974E-3 | 0.866E-3 | 1.105E-3 | |

| | | | | | | | | | |
|---|---|---|---|---|---|---|---|---|---|
| 7 days old on both diets | D | 1 | 0.0761 | 26.61 | <0.0001 | Fixed | Fixed | Fixed | 0.3081 |
| | L | 97 | 7.572E-3 | 2.63 | <0.0001 | 0.304E-3 | 0.156E-3 | 0.452E-3 | |
| | D×L | 97 | 2.88E-3 | 2.20 | <0.0001 | 0.204E-3 | 0.092E-3 | 0.315E-3 | |
| | Rep(D×L) | 339 | 1.288 | 1.13 | 0.0823 | 0.054E-3 | -0.02E-3 | 0.131E-3 | |
| | ϵ | 1026 | 1.141E-3 | - | - | 1.141E-3 | 1.049E-3 | 1.247E-3 | |
| 16 days old on control diet | L | 92 | 3.580E-3 | 3.35 | <0.0001 | 0.325E-3 | 0.187E-3 | 0.464E-3 | 0.2395 |
| | Rep(L) | 164 | 1.066E-3 | 1.03 | 0.3926 | 0.012E-3 | -0.08E-3 | 0.102E-3 | |
| | ϵ | 483 | 1.032E-3 | - | - | 1.032E-3 | 0.914E-3 | 1.176E-3 | |
| 16 days old on restricted diet | L | 92 | 4.369E-3 | 3.69 | <0.0001 | 0.419E-3 | 0.250E-3 | 0.588E-3 | 0.2779 |
| | Rep(L) | 162 | 1.173E-3 | 1.08 | 0.2741 | 0.031E-3 | -0.07E-3 | 0.132E-3 | |
| | ϵ | 485 | 1.089E-3 | - | - | 1.089E-3 | 0.964E-3 | 1.241E-3 | |
| 16 days old on both diets | D | 1 | 0.0560 | 31.80 | <0.0001 | Fixed | Fixed | Fixed | 0.2580 |
| | L | 92 | 6.126E-3 | 3.46 | <0.0001 | 0.285E-3 | 0.164E-3 | 0.406E-3 | |
| | D×L | 92 | 1.770E-3 | 1.57 | 0.0030 | 0.084E-3 | 0.014E-3 | 0.154E-3 | |
| | Rep(D×L) | 326 | 1.119E-3 | 1.05 | 0.2718 | 0.022E-3 | -0.04E-3 | 0.088E-3 | |
| | ϵ | 968 | 1.061E-3 | - | - | 1.061E-3 | 0.972E-3 | 1.162E-3 | |
| 7 and 16 days old on control diet | A | 1 | 1.080E-3 | 0.50 | 0.4801 | Fixed | Fixed | Fixed | 0.3095 |
| | L | 92 | 8.166E-3 | 3.80 | <0.0001 | 0.391E-3 | 0.230E-3 | 0.552E-3 | |
| | A×L | 92 | 2.151E-3 | 1.86 | <0.0001 | 0.129E-3 | 0.045E-3 | 0.214E-3 | |
| | Rep(A×L) | 329 | 1.156E-3 | 1.00 | 0.5109 | -1.63E-6 | -0.07E-3 | 0.066E-3 | |
| | ϵ | 964 | 1.160E-3 | - | - | 1.160E-3 | 1.063E-3 | 1.271E-3 | |
| 7 and 16 days old on restricted diet | A | 1 | 2.790E-3 | 1.28 | 0.2610 | Fixed | Fixed | Fixed | 0.2588 |
| | L | 92 | 5.845E-3 | 2.65 | <0.0001 | 0.238E-3 | 0.120E-3 | 0.356E-3 | |
| | A×L | 92 | 2.203E-3 | 1.75 | 0.0003 | 0.123E-3 | 0.040E-3 | 0.207E-3 | |
| | Rep(A×L) | 326 | 1.234E-3 | 1.19 | 0.0233 | 0.074E-3 | -3.67E-6 | 0.151E-3 | |
| | ϵ | 971 | 1.034E-3 | - | - | 1.034E-3 | 0.948E-3 | 1.133E-3 | |
| 7 and 16 days old on both diets | A | 1 | 3.387E-3 | 1.62 | 0.2062 | Fixed | Fixed | Fixed | 0.2871 |
| | D | 1 | 0.1252 | 52.12 | <0.0001 | Fixed | Fixed | Fixed | |
| | L | 92 | 0.0116 | 5.18 | <0.0001 | 0.304E-3 | 0.188E-3 | 0.420E-3 | |
| | A×D | 1 | 0.191E-3 | 0.08 | 0.7723 | Fixed | Fixed | Fixed | |
| | A×L | 92 | 2.096E-3 | 0.92 | 0.6517 | -0.01E-3 | -0.07E-3 | 0.046E-3 | |
| | D×L | 92 | 2.410E-3 | 1.06 | 0.3912 | 8.824E-6 | -0.05E-3 | 0.071E-3 | |
| | A×D×L | 92 | 2.275E-3 | 1.89 | <0.0001 | 0.139E-3 | 0.051E-3 | 0.228E-3 | |
| | Rep(A×D×L) | 655 | 1.195E-3 | 1.09 | 0.0881 | 0.036E-3 | -0.02E-3 | 0.087E-3 | |
| | ϵ | 1935 | 1.097E-3 | - | - | 1.097E-3 | 1.031E-3 | 1.170E-3 | |
| 25 days old on control diet | L | 84 | 3.944E-3 | 2.92 | <0.0001 | 0.316E-3 | 0.163E-3 | 0.469E-3 | 0.2631 |
| | Rep(L) | 143 | 1.324E-3 | 1.50 | 0.0009 | 0.146E-3 | 0.037E-3 | 0.254E-3 | |
| | ϵ | 490 | 0.885E-3 | - | - | 0.885E-3 | 0.783E-3 | 1.007E-3 | |
| 25 days old on restricted diet | L | 84 | 3.177E-3 | 2.14 | <0.0001 | 0.232E-3 | 0.091E-3 | 0.373E-3 | 0.1476 |
| | Rep(L) | 124 | 1.482E-3 | 1.11 | 0.2319 | 0.048E-3 | -0.09E-3 | 0.186E-3 | |
| | ϵ | 433 | 1.340E-3 | - | - | 1.340E-3 | 1.178E-3 | 1.538E-3 | |
| 25 days old on both diets | D | 1 | 0.0364 | 23.27 | <0.0001 | Fixed | Fixed | Fixed | 0.1985 |
| | L | 84 | 5.166E-3 | 3.12 | <0.0001 | 0.237E-3 | 0.125E-3 | 0.350E-3 | |
| | D×L | 84 | 1.654E-3 | 1.18 | 0.1608 | 0.035E-3 | -0.04E-3 | 0.113E-3 | |
| | Rep(A×D×L) | 267 | 1.397E-3 | 1.27 | 0.0058 | 0.100E-3 | 0.014E-3 | 0.186E-3 | |
| | ϵ | 923 | 1.098E-3 | - | - | 1.098E-3 | 1.005E-3 | 1.206E-3 | |
| 7, 16 and 25 days old on both diets | A | 2 | 1.036E-3 | 0.81 | 0.4453 | Fixed | Fixed | Fixed | 0.9763 |
| | D | 1 | 0.063E-9 | 0.00 | 0.9998 | Fixed | Fixed | Fixed | |
| | L | 82 | 0.0136 | -0.01 | - | 0.0247 | 0.0234 | 0.0260 | |
| | A×D | 2 | 0.526E-3 | 0.00 | 0.9991 | Fixed | Fixed | Fixed | |
| | A×L | 164 | 2.111E-3 | 0.00 | 1.0000 | -0.0734 | -0.0772 | -0.0696 | |
| | D×L | 82 | 2.400E-3 | 0.00 | 1.0000 | -0.0489 | -0.0514 | -0.0464 | |
| | A×D×L | 164 | 2.0827 | 1649.96 | <0.0001 | 0.1467 | 0.1391 | 0.1543 | |
| | Rep(A×D×L) | 859 | 1.252E-3 | 1.12 | 0.0208 | 0.047E-3 | 4.1E-6 | 0.091E-3 | |
| | ϵ | 2636 | 1.119E-3 | - | - | 1.119E-3 | 1.061E-3 | 1.182E-3 | |

**Table S8.** Summary of full and reduced ANOVA models for balanced data assessing locomotor activity of DGRP flies at 7, 16 and 25 days old and fed control and restricted diets. Diet and age were considered fixed effects, and DGRP line is a random effect.

| Analysis | Source | DF | MS | F Value | *p*-value | $\sigma^2$ | Lower | Upper | $H^2$ |
|---|---|---|---|---|---|---|---|---|---|
| 7 days old on control diet | L | 97 | 3.7961 | 5.21 | <0.0001 | 0.4034 | 0.2627 | 0.5442 | 0.3717 |
| | Rep(L) | 170 | 0.7225 | 1.06 | 0.3143 | 0.0152 | -0.0505 | 0.0809 | |

| | | | | | | | | | |
|---|---|---|---|---|---|---|---|---|---|
| | ϵ | 502 | 0.6819 | - | - | 0.6819 | 0.6048 | 0.7748 | |
| 7 days old on restricted diet | L | 97 | 5.3835 | 5.11 | <0.0001 | 0.5805 | 0.3756 | 0.7855 | 0.4221 |
| | Rep(L) | 168 | 1.0182 | 1.28 | 0.0219 | 0.0848 | -0.0055 | 0.1750 | |
| | ϵ | 492 | 0.7949 | - | - | 0.7949 | 0.7042 | 0.9044 | |
| 7 days old on both diets | D | 1 | 0.0010 | 0.00 | 0.9807 | Fixed | Fixed | Fixed | 0.4004 |
| | L | 97 | 7.4081 | 4.20 | <0.0001 | 0.3762 | 0.2328 | 0.5197 | |
| | D×L | 97 | 1.7623 | 1.98 | <0.0001 | 0.1164 | 0.0488 | 0.1841 | |
| | Rep(D×L) | 338 | 0.8695 | 1.18 | 0.0299 | 0.0496 | -0.0051 | 0.1043 | |
| | ϵ | 994 | 0.7378 | - | - | 0.7378 | 0.6770 | 0.8073 | |
| 16 days old on control diet | L | 90 | 5.8590 | 4.64 | <0.0001 | 0.7390 | 0.4590 | 1.0190 | 0.4250 |
| | Rep(L) | 141 | 1.2394 | 1.24 | 0.0575 | 0.0998 | -0.0366 | 0.2361 | |
| | ϵ | 370 | 1.0000 | - | - | 1.0000 | 0.8702 | 1.1614 | |
| 16 days old on restricted diet | L | 90 | 4.8761 | 5.74 | <0.0001 | 0.5939 | 0.3863 | 0.8014 | 0.3942 |
| | Rep(L) | 156 | 0.8563 | 0.94 | 0.6761 | -0.0231 | -0.1077 | 0.0616 | |
| | ϵ | 413 | 0.9127 | - | - | 0.9127 | 0.8000 | 1.0512 | |
| 16 days old on both diets | D | 1 | 34.0410 | 13.64 | 0.0004 | Fixed | Fixed | Fixed | 0.4126 |
| | L | 90 | 7.8453 | 2.93 | <0.0001 | 0.4108 | 0.2248 | 0.5969 | |
| | D×L | 90 | 2.6653 | 2.55 | <0.0001 | 0.2593 | 0.1307 | 0.3878 | |
| | Rep(D×L) | 297 | 1.0382 | 1.09 | 0.1848 | 0.0348 | -0.0435 | 0.1130 | |
| | ϵ | 783 | 0.9540 | - | - | 0.9540 | 0.8661 | 1.0560 | |
| 7 and 16 days old on control diet | A | 1 | 18.4781 | 5.14 | 0.0258 | Fixed | Fixed | Fixed | 0.4179 |
| | L | 90 | 6.0150 | 1.56 | 0.0186 | 0.1604 | 0.0066 | 0.3141 | |
| | A×L | 90 | 3.8375 | 3.90 | <0.0001 | 0.4286 | 0.2576 | 0.5996 | |
| | Rep(A×L) | 302 | 0.9706 | 1.18 | 0.0354 | 0.0590 | -0.0123 | 0.1303 | |
| | ϵ | 836 | 0.8204 | - | - | 0.8204 | 0.7471 | 0.9051 | |
| 7 and 16 days old on restricted diet | A | 1 | 2.0177 | 1.01 | 0.3168 | Fixed | Fixed | Fixed | 0.4073 |
| | L | 90 | 8.0030 | 3.91 | <0.0001 | 0.4236 | 0.2510 | 0.5961 | |
| | A×L | 90 | 2.0448 | 2.23 | <0.0001 | 0.1603 | 0.0766 | 0.2440 | |
| | Rep(A×L) | 315 | 0.9114 | 1.07 | 0.2198 | 0.0243 | -0.0357 | 0.0842 | |
| | ϵ | 870 | 0.8496 | - | - | 0.8496 | 0.7751 | 0.9354 | |
| 7 and 16 days old on both diets | A | 1 | 4.1015 | 1.26 | 0.2641 | Fixed | Fixed | Fixed | 0.4140 |
| | D | 1 | 21.2443 | 11.45 | 0.0010 | Fixed | Fixed | Fixed | |
| | L | 90 | 11.8439 | 4.38 | <0.0001 | 0.3387 | 0.2019 | 0.4755 | |
| | A×D | 1 | 17.4111 | 7.00 | 0.0096 | Fixed | Fixed | Fixed | |
| | A×L | 90 | 3.3723 | 1.31 | 0.1044 | 0.0586 | -0.0314 | 0.1486 | |
| | D×L | 90 | 1.9098 | 0.74 | 0.9235 | -0.0501 | -0.1200 | 0.0197 | |
| | A×D×L | 90 | 2.5778 | 2.72 | <0.0001 | 0.2429 | 0.1271 | 0.3586 | |
| | Rep(A×D×L) | 617 | 0.9404 | 1.13 | 0.0352 | 0.0413 | -0.0056 | 0.0880 | |
| | ϵ | 1706 | 0.8353 | - | - | 0.8353 | 0.7820 | 0.8943 | |
| 25 days old on control diet | L | 82 | 5.9119 | 3.76 | <0.0001 | 0.6601 | 0.3753 | 0.9448 | 0.4154 |
| | Rep(L) | 126 | 1.5622 | 1.68 | <0.0001 | 0.2395 | 0.0919 | 0.3872 | |
| | ϵ | 370 | 0.9290 | - | - | 0.9290 | 0.8084 | 1.0788 | |
| 25 days old on restricted diet | L | 82 | 2.9789 | 2.78 | <0.0001 | 0.3083 | 0.1512 | 0.4654 | 0.2404 |
| | Rep(L) | 115 | 1.0713 | 1.10 | 0.2566 | 0.0372 | -0.0761 | 0.1504 | |
| | ϵ | 343 | 0.9741 | - | - | 0.9741 | 0.8433 | 1.1380 | |
| 25 days old on both diets | D | 1 | 0.9845 | 0.55 | 0.4582 | Fixed | Fixed | Fixed | 0.3436 |
| | L | 82 | 6.5560 | 3.32 | <0.0001 | 0.3859 | 0.2017 | 0.5701 | |
| | D×L | 82 | 1.9615 | 1.50 | 0.0093 | 0.1118 | -0.0045 | 0.2281 | |
| | Rep(A×D×L) | 241 | 1.3279 | 1.40 | 0.0005 | 0.1434 | 0.0519 | 0.2350 | |
| | ϵ | 713 | 0.9507 | - | - | 0.9507 | 0.8592 | 1.0576 | |
| 7, 16 and 25 days old on both diets | A | 2 | 18.2214 | 5.81 | 0.0037 | Fixed | Fixed | Fixed | 0.3782 |
| | D | 1 | 16.7769 | 7.87 | 0.0062 | Fixed | Fixed | Fixed | |
| | L | 79 | 14.3821 | 4.28 | <0.0001 | 0.2863 | 0.1684 | 0.4041 | |
| | A×D | 2 | 5.3980 | 2.64 | 0.0747 | Fixed | Fixed | Fixed | |
| | A×L | 158 | 3.3132 | 1.54 | 0.0037 | 0.0881 | 0.0208 | 0.1553 | |
| | D×L | 79 | 2.2244 | 1.05 | 0.3943 | 0.0054 | -0.0380 | 0.0488 | |
| | A×D×L | 158 | 2.1464 | 2.00 | <0.0001 | 0.1647 | 0.0886 | 0.2408 | |
| | Rep(A×D×L) | 796 | 1.0652 | 1.19 | 0.0012 | 0.0661 | 0.0221 | 0.1100 | |
| | ϵ | 2228 | 0.8950 | - | - | 0.8950 | 0.8447 | 0.9500 | |

**Table S9.** Summary of full and reduced ANOVA models for balanced data assessing HKDT of DGRP flies at 7, 16 and 25 days old and fed control and restricted diets. Diet and age were considered fixed effects, and DGRP line is a random effect.

| Analysis | Source | DF | MS | F Value | *p*-value | $\sigma^2$ | Lower | Upper | $H^2$ |
|---|---|---|---|---|---|---|---|---|---|

| | | | | | | | | | |
|---|---|---|---|---|---|---|---|---|---|
| 7 days old on control diet | L | 93 | 490.7827 | 3.85 | <0.0001 | 53.1426 | 31.8696 | 74.4157 | 0.3636 |
| | Rep(L) | 151 | 123.3621 | 1.33 | 0.0150 | 12.0271 | -1.7060 | 25.7602 | |
| | $\epsilon$ | 424 | 93.0129 | - | - | 93.0129 | 81.6604 | 106.92 | |
| 7 days old on restricted diet | L | 93 | 394.1051 | 3.60 | <0.0001 | 51.3928 | 28.8399 | 73.9457 | 0.3264 |
| | Rep(L) | 138 | 109.0432 | 1.03 | 0.4159 | 1.3612 | -13.0343 | 15.7566 | |
| | $\epsilon$ | 325 | 106.0603 | - | - | 106.06 | 91.4711 | 124.46 | |
| 7 days old on both diets | D | 1 | 16006 | 58.96 | <0.0001 | Fixed | Fixed | Fixed | 0.3475 |
| | L | 93 | 569.9046 | 1.94 | 0.0008 | 23.0790 | 7.6422 | 38.5158 | |
| | D×L | 93 | 292.9536 | 2.48 | <0.0001 | 29.4762 | 15.1484 | 43.8040 | |
| | Rep(D×L) | 289 | 116.5247 | 1.18 | 0.0417 | 7.5481 | -2.3807 | 17.4769 | |
| | $\epsilon$ | 749 | 98.6743 | - | - | 98.6743 | 89.3964 | 109.48 | |
| 16 days old on control diet | L | 58 | 231.8209 | 1.38 | 0.1179 | 18.8209 | -14.1136 | 51.7554 | 0.1188 |
| | Rep(L) | 55 | 168.2373 | 1.21 | 0.2082 | 16.0936 | -35.9378 | 68.1249 | |
| | $\epsilon$ | 100 | 139.6117 | - | - | 139.61 | 107.76 | 188.10 | |
| 16 days old on restricted diet | L | 58 | 267.0131 | 3.02 | <0.0001 | 35.4535 | 12.4704 | 58.4367 | 0.3224 |
| | Rep(L) | 83 | 86.7528 | 1.16 | 0.2004 | 6.0470 | -14.2512 | 26.3451 | |
| | $\epsilon$ | 180 | 74.4996 | - | - | 74.4996 | 61.2202 | 92.6476 | |
| 16 days old on both diets | D | 1 | 1470.7298 | 7.25 | 0.0088 | Fixed | Fixed | Fixed | 0.2484 |
| | L | 58 | 247.4582 | 1.06 | 0.4117 | 1.9382 | -17.3029 | 21.1793 | |
| | D×L | 58 | 227.3544 | 1.92 | 0.0010 | 30.3665 | 0.6431 | 60.0899 | |
| | Rep(D×L) | 138 | 119.2285 | 1.22 | 0.0839 | 11.1404 | -11.4137 | 33.6944 | |
| | $\epsilon$ | 280 | 97.7539 | - | - | 97.7539 | 83.3861 | 116.21 | |
| 7 and 16 days old on control diet | A | 1 | 32572 | 162.97 | <0.0001 | Fixed | Fixed | Fixed | 0.2540 |
| | L | 57 | 341.5793 | 1.49 | 0.0724 | 12.8513 | -5.1703 | 30.8729 | |
| | A×L | 57 | 222.6583 | 1.62 | 0.0091 | 20.7474 | -2.3855 | 43.8802 | |
| | Rep(A×L) | 152 | 140.9800 | 1.43 | 0.0034 | 18.6465 | -0.0421 | 37.3351 | |
| | $\epsilon$ | 377 | 98.6855 | - | - | 98.6855 | 85.9845 | 114.44 | |
| 7 and 16 days old on restricted diet | A | 1 | 6631.1441 | 33.53 | <0.0001 | Fixed | Fixed | Fixed | 0.3503 |
| | L | 57 | 431.9360 | 2.00 | 0.0050 | 20.4265 | 2.5801 | 38.2729 | |
| | A×L | 57 | 214.0833 | 2.39 | <0.0001 | 23.7360 | 7.0134 | 40.4587 | |
| | Rep(A×L) | 172 | 89.0259 | 1.09 | 0.2514 | 3.3026 | -8.5758 | 15.1810 | |
| | $\epsilon$ | 407 | 81.8909 | - | - | 81.8909 | 71.7092 | 94.4185 | |
| 7 and 16 days old on both diets | A | 1 | 36098 | 187.00 | <0.0001 | Fixed | Fixed | Fixed | 0.3000 |
| | D | 1 | 253.0245 | 0.95 | 0.3323 | Fixed | Fixed | Fixed | |
| | L | 57 | 426.9033 | 1.45 | 0.1454 | 7.3664 | -5.5699 | 20.3027 | |
| | A×D | 1 | 7162.0576 | 36.83 | <0.0001 | Fixed | Fixed | Fixed | |
| | A×L | 57 | 209.3845 | 0.98 | 0.5233 | -0.3679 | -14.0096 | 13.2737 | |
| | D×L | 57 | 294.8821 | 1.38 | 0.1186 | 8.9778 | -6.7782 | 24.7339 | |
| | A×D×L | 57 | 210.4926 | 1.87 | 0.0004 | 22.5833 | 3.7607 | 41.4059 | |
| | Rep(A×D×L) | 324 | 113.3995 | 1.26 | 0.0057 | 10.5982 | -0.0089 | 21.2054 | |
| | $\epsilon$ | 784 | 89.9669 | - | - | 89.9669 | 81.6851 | 99.5820 | |
| 25 days old on control diet | L | 58 | 266.8686 | 2.47 | 0.0002 | 39.8890 | 14.5473 | 65.2308 | 0.2098 |
| | Rep(L) | 61 | 107.0241 | 0.71 | 0.9318 | -21.3864 | -40.1835 | -2.5894 | |
| | $\epsilon$ | 137 | 150.2157 | - | - | 150.22 | 120.14 | 193.25 | |
| 25 days old on restricted diet | L | 58 | 218.8378 | 1.06 | 0.4060 | 3.0713 | -16.0853 | 22.2279 | 0.0430 |
| | Rep(L) | 65 | 201.4163 | 2.94 | <0.0001 | 67.1812 | 27.8347 | 106.53 | |
| | $\epsilon$ | 137 | 68.4057 | - | - | 68.4057 | 54.7105 | 88.0039 | |
| 25 days old on both diets | D | 1 | 414.8376 | 1.77 | 0.1874 | Fixed | Fixed | Fixed | 0.1737 |
| | L | 58 | 214.3308 | 0.80 | 0.7957 | -7.0070 | -23.1949 | 9.1808 | |
| | D×L | 58 | 262.7501 | 1.71 | 0.0058 | 29.9785 | 3.7040 | 56.2530 | |
| | Rep(A×D×L) | 126 | 155.7185 | 1.42 | 0.0085 | 23.2143 | -0.1832 | 46.6118 | |
| | $\epsilon$ | 274 | 109.3107 | - | - | 109.31 | 93.0895 | 130.20 | |
| 7, 16 and 25 days old on both diets | A | 2 | 21254 | 111.19 | <0.0001 | Fixed | Fixed | Fixed | 0.2711 |
| | D | 1 | 27.2007 | 0.08 | 0.7829 | Fixed | Fixed | Fixed | |
| | L | 43 | 395.7983 | 1.04 | 0.4640 | 0.5485 | -9.9679 | 11.0649 | |
| | A×D | 2 | 3430.1651 | 16.71 | <0.0001 | Fixed | Fixed | Fixed | |
| | A×L | 86 | 206.4383 | 0.93 | 0.6400 | -1.8587 | -13.0767 | 9.3592 | |
| | D×L | 43 | 395.9592 | 1.82 | 0.0092 | 13.8909 | -0.7253 | 28.5071 | |
| | A×D×L | 86 | 220.9922 | 1.85 | <0.0001 | 22.9778 | 5.8859 | 40.0697 | |
| | Rep(A×D×L) | 359 | 120.7862 | 1.26 | 0.0038 | 11.4927 | 1.3147 | 21.6706 | |
| | $\epsilon$ | 838 | 95.5975 | - | - | 95.5975 | 87.0658 | 105.45 | |

**Table S11.** Summary of full and reduced ANOVA models assessing lifespan of DGRP flies fed control and restricted diets. Diet and age were considered fixed effects, and DGRP line is a random effect.

| Analysis | Source | DF | MS | F Value | *p*-value | $\sigma^2$ | Lower | Upper | $H^2$ |
|---|---|---|---|---|---|---|---|---|---|
| Control diet | L | 95 | 7601.7568 | 13.39 | <0.0001 | 43.6137 | 29.4308 | 57.7966 | 0.3909 |
| | Rep(L) | 171 | 600.3582 | 8.83 | <0.0001 | 8.8832 | 6.8248 | 10.9416 | |
| | $\epsilon$ | 15637 | 67.9693 | - | - | 67.9693 | 66.4875 | 69.5014 | |
| Restricted diet | L | 97 | 7401.8553 | 14.06 | <0.0001 | 35.5377 | 24.5383 | 46.5371 | 0.5319 |
| | Rep(L) | 171 | 565.1848 | 18.07 | <0.0001 | 7.3473 | 5.6598 | 9.0349 | |
| | $\epsilon$ | 18894 | 31.2792 | - | - | 31.2792 | 30.6579 | 31.9197 | |
| Both diets | D | 1 | 21019 | 8.65 | 0.0041 | Fixed | Fixed | Fixed | 0.4525 |
| | L | 97 | 11589 | 3.79 | <0.0001 | 24.8539 | 14.5188 | 35.1891 | |
| | D×L | 95 | 3120.5366 | 5.77 | <0.0001 | 14.7233 | 9.4495 | 19.9971 | |
| | Rep(D×L) | 342 | 582.7715 | 12.17 | <0.0001 | 8.0676 | 6.7788 | 9.3565 | |
| | $\epsilon$ | 34531 | 47.8939 | - | - | 47.8939 | 47.1875 | 48.6164 | |

**Table S12.** Summary of the Cox Proportional Hazards model assessing lifespan of DGRP flies fed control and restricted diets.

| Source | β | se(β) | z | *p*-value |
|---|---|---|---|---|
| Diet | 0.25859 | 0.01117 | 23.14 | <0.0001 |
